# isotracer: An R package for the analysis of tracer addition experiments

**DOI:** 10.1101/2021.08.09.455668

**Authors:** Matthieu Bruneaux, Andrés López-Sepulcre

## Abstract

1. Tracer addition experiments, particularly using isotopic tracers, are becoming increasingly important in a variety of studies aiming at characterizing the flows of molecules or nutrients at different levels of biological organization, from the cellular and tissue levels, to the organismal and ecosystem levels.
2. We present an approach based on Hidden Markov Models (HMM) to estimate nutrient flow parameters across a network, and its implementation in the R package isotracer.
3. The isotracer package is capable of handling a variety of tracer study designs, including continuous tracer drips, pulse experiments, and pulse-chase experiments. It can also take into account tracer decay when radioactive isotopes are used.
4. To illustrate its use, we present three case studies based on published data and spanning different levels of biological organization: a molecular-level study of protein synthesis and degradation in *Arabidopsis thaliana*, an organismal-level study of phosphorus incorporation in the eelgrass *Zostera marina*, and an ecosystem-level study of nitrogen dynamics in Trinidadian montane streams.
5. With these case studies, we illustrate how isotracer can be used to estimate uptake rates, turnover rates, and total flows, as well as their uncertainty. We also show how to perform model selection to compare alternative hypotheses.
6. We conclude by discussing isotracer’s further applications, limitations, and possible future improvements and expansions.

## 1 Introduction

Tracer addition experiments are an increasingly common tool used to answer a wide variety of biological, ecological and evolutionary questions. These experiments consist in injecting a labelled element into a biological system and tracing its fate throughout the system compartments across time to estimate flows of material across the compartments. Their applications encompass all levels of biological organization, from cells and tissues (Yuan et al., 2006; Allen and Young, 2020) to organisms (Kim et al., 2016; Williams et al., 2016), communities (Shik et al., 2018; Freeman et al., 2013), and ecosystems (Riis et al., 2012; Collins et al., 2016). Tracer additions have a long history. They have been used in seminal experiments revealing the nature of the genetic material (Hershey and Chase, 1952), and more recently in high-throughput metabolomic techniques (e.g. Yuan et al. (2006); Li et al. (2017)). They are also becoming a common tool to study the dynamics of nutrients or elements in a variety of ecosystems including: water dynamics in a savannah (Kulmatiski et al., 2010), carbon in coral reefs (de Goeij et al., 2013), nitrogen in decidious forest soil (Goodale et al., 2015), the effects of reindeer urine in tundra ecosystems, (Barthelemy et al., 2018), and nutrient cycling in a variety of aquatic environments (Sánchez-Carrillo and Álvarez Cobelas, 2017). However, gaining reliable quantitative insights and making robust inferences from such data requires appropriate statistical techniques which can sometimes be challenging to develop and use.

Analysis of tracer addition data typically requires models incorporating a time component via a system of differential equations describing the flows of material across connected compartments, which can be outside the typical statistical expertise of biologists or field geologists. While some sophisticated frameworks exist to analyze pharmacokinetics data (Gelman et al. (1996), Lunn et al. (2002) for PKBugs, Gillespie et al. (2021) for Torsten, currently in development) or metabolomic data (Weindl et al., 2015), other disciplines such as evolutionary biology, ecology or ecosystem science suffer from a lack of a standard analytical tool to make statistically rigorous inferences from tracer additions. For example, analysis of tracer addition experiments to study food web dynamics is usually done by fitting mass balance equations focusing on one trophic compartment at a time, and trophic relationships are assumed to be known a priori, including the relative proportions of diet sources when a consumer feeds on more than one source (Dodds et al., 2000; Mulholland et al., 2000; Collins et al., 2016). Such an approach does not take into account the uncertainty in assumed trophic relationships (Ainsworth et al., 2010; Dodds et al., 2014), and does not allow either the uncertainty in flow estimates at one trophic level to be taken into account when estimating flow in the rest of the trophic network. Statistically reliable estimates of parameter uncertainty are particularly important when performing comparative or experimental studies (e.g. Whiles et al. (2013); Collins et al. (2016); Norman et al. (2017); Tank et al. (2018)).

We developed the isotracer R package in an effort to make a statistically rigorous framework for such analyses more easily accessible for researchers. Our package implements the mathematical framework recently described in López-Sepulcre et al. (2020). It considers a whole network system simultaneously and uses a Bayesian approach to incorporate experimental uncertainty into model parameter estimates. For a given set of parameter values, likelihood is calculated by solving numerically the system of differential equations governing the material flows, and by comparing the expected compartment sizes and tracer contents to observations. The current version of isotracer implements first-order kinetics for material transfer across compartment, allows replicated units in an experimental design and can take into account categorical covariates to model treatment effects. By using a Bayesian framework, we can make statistically rigorous inferences, estimate parameter uncertainty and calculate derived parameters such as network-wide properties (e.g. total nutrient flow in an ecosystem). Model comparison is possible when several plausible models exist. Additionally, the package also allows researchers to perform power analyses when designing their tracer addition experiments in order to make the most out of such cost- and effort-intensive studies.

We first describe the mathematical framework used in our modelling approach and an overview of its implementation in the package. We then illustrate the use of the package through three case studies based on published datasets: protein turnover in *Arabidopsis thaliana* leaves (Li et al., 2017), phosphate uptake in *Zostera marina* individuals (McRoy and Barsdate, 1970), and nitrogen flows in a Trinidadian mountain stream (Collins et al., 2016). Detailed tutorials to reproduce those case studies are available as an appendix.

## 2 Mathematical framework

The system of interest can be represented as a network of connected compartments (Figure 1). As examplified in our case studies, compartments can be anything from pools of molecules (e.g. intermediates in cellular biosynthesis pathways), to tissues (e.g. blood, liver, and muscle; or leaf, stem, and root), to species or functional groups (e.g. algae, invertebrate grazer, fish predator). Each compartment represents a distinct pool of matter of any given chemical element of interest, and that matter can flow from one compartment to another when compartments are connected. In a tracer addition experiment, a very small fraction of labelled or marked element of interest (the tracer) is added. The *marked* fraction is followed in time throughout the system’s compartment in comparison to the natural *unmarked* fraction. In essence, modelling data from a tracer addition experiment consists of comparing the expected trajectories of tracer with observations, given a set of parameter values. In the case of stable isotope studies, the tracer is often a naturally rare heavy isotopic form of the element of interest (e.g. ^2^H, ^13^C, ^15^N, ^18^O, ^34^S) which is used to enrich the pools of the corresponding naturally abundant unmarked form (^1^H, ^12^C, ^14^N, ^16^O, ^32^S). Observations usually comprise both the total pool size for each compartment (i.e. the sum of the quantities of heavy and light isotopes) and the proportion of tracer in each pool (i.e. the proportion of heavy isotope to the total amount of heavy and light isotopes, often measured as *δ* enrichment values (Fry, 2006)). In the case where only the tracer is being tracked (e.g. a radioactive tracer such as ^32^P), the general framework presented above can be used by considering that the observed sizes represent the tracer alone, and that the observed marked proportions are always one. Radioactive decay is taken into account in the turnover rate of the tracer. It is important to note, however, that the method works for any traceable marker (e.g. immunolabelled molecules), not just rare isotopes.

**Figure 1:**
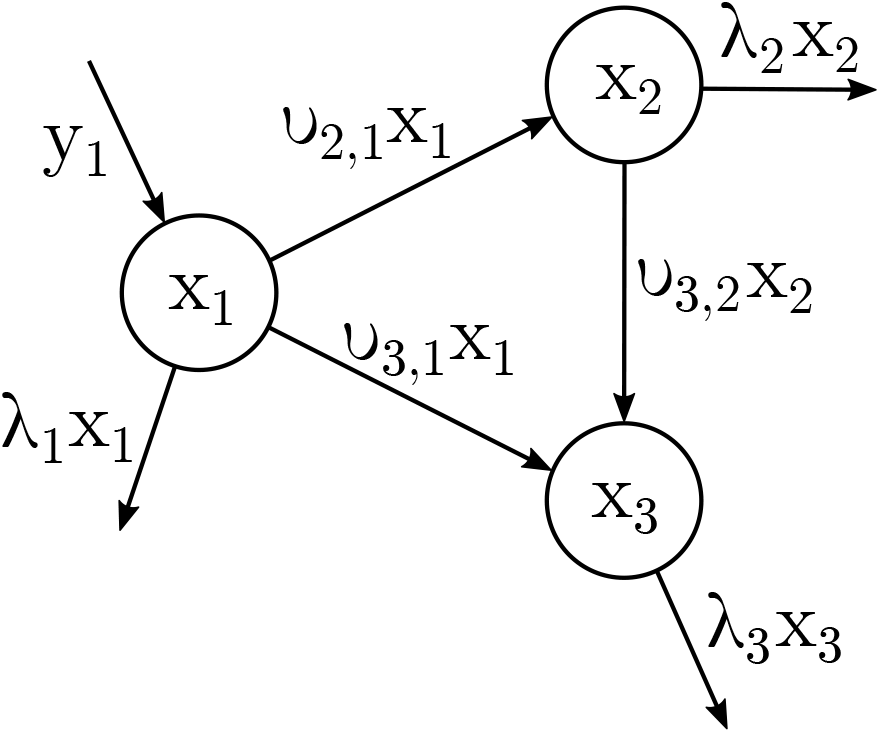
Example of a basic network with three compartments. The *x*_*i*_ values in each compartment are quantities of matter. The values along arrows are rates or flows of matter (quantity per time unit). *y*_1_ is a rate (quantity per time unit); *υ*_*i,j*_ and *λ*_*i*_ are rate coefficients of first-order kinetics (per time unit). *x*_*i*_ and *y*_1_ are functions of time *t* while *υ*_*i,j*_ and *λ*_*i*_ are constant rate coefficients.

The mathematical model used to calculate the likelihood of a set of parameter values is detailed in the sections below, and can be divided into two parts. The first part is the calculation of the expected movement of material across the network for those parameter values, which allows to predict the latent state of the network at any point in time. The second part is the likelihood calculation through the incorporation of process and observation error: observations are assumed to be generated via some statistical distribution parameterized with the expected latent states of the network calculated previously.

### 2.1 General formulation of transfer equations

The mathematical framework described here is similar to the one described in López-Sepulcre et al. (2020). We try to keep mathematical notations consistent with it as much as possible. While the presentation in López-Sepulcre et al. (2020) focuses on discrete-time modelling, we present only the equivalent continuous-time approach here, which allows to specify the model based on a system of differential equations.

Let’s consider a network with *N* distinct compartments (*N* = 3 in Figure 1). Each compartment is a pool of the element of interes (e.g. sulphur), and we can define the network state at time *t* by a *N* × 1 vector 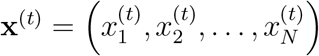 where 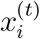 indicates the quantity of material in compartment *i* at time *t*. Flows between connected compartments are assumed to follow first-order kinetics, and the transfer rate coefficients are contained in a matrix **ϒ** (capital upsilon) where each coefficient *υ*_*i,j*_ determines the transfer rate from compartment *j* to *i* as 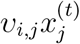. Transfer rates can represent a variety of processes depending on the modelled system, including: chemical transformations, resource allocation among tissues, nutrient uptake, or consumption among trophic levels. In addition to flowing between compartments, matter can be lost from the system (e.g. through excretion or emigration): each compartment *i* has a loss rate coefficient *λ*_*i*_ which also defines first-order kinetics for the material exiting the system from this compartment. Finally, some exogenous input can add matter to the system, and is defined for each compartment *i* as an input function 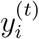 giving the input rate for this compartment. Exogenous inputs can represent a variety of scenarios such as nutrients entering an ecosystem from adjacent systems (e.g. upstream in rivers, terrestrial or aerial inputs in aquatic ecosystems), available food or nutrients to be eaten or uptaken by animals or plants, or pollutants in an organism or ecosystem, to name a few examples.

The evolution of such a network system is entirely described by solving for **x**^(*t*)^ in a linear system of first-order differential equations, for a given 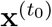 defining the initial conditions. For the simple network example shown in Figure 1, the system of equations is:

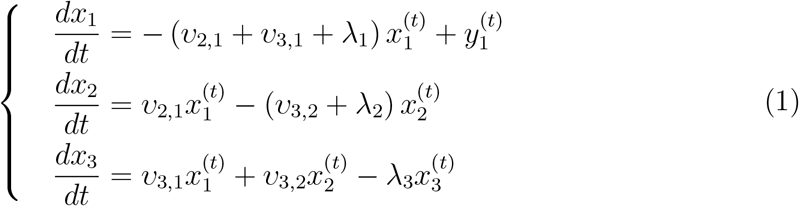

which can be written in a matrix form:

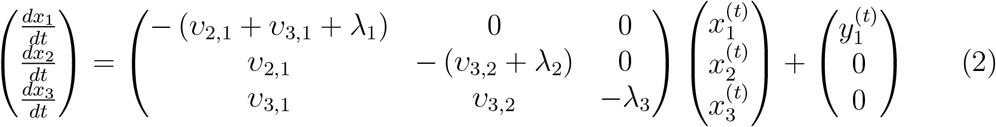

In this form, non-zero *υ*_*i,j*_ outside the diagonal of the matrix defines a transfer from *j* to *i*, while the diagonal coefficients are the overall turnover rate coefficients for each compartment. The added vector **y**^(*t*)^ represents the exogenous input rates.

In the more general case, the system of differential equations can be written:

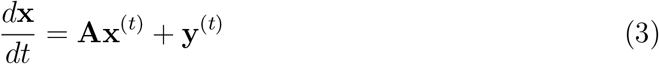

with transfer matrix **A** = **ϒ** − **K** · **I**_*N*_ where **K** is a vector containing the turnover rate coefficients for each compartments such that, for compartment 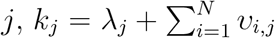, and **I**_*N*_ is the identity matrix.

### 2.2 Likelihood calculation

As mentioned above, tracer addition data usually comprise time series of the compartment *sizes* as well as their *labelled proportions*. We use interchangeably the terms “compartments” and “pools”, and the terms “tracer”, “labelled” element and “marked” element to refer to the tracer being added, and “unlabelled” or “unmarked” element for the corresponding common isotope being measured at the same time as the tracer.

To model compartment sizes and labelled proportions, it is necessary to distinguish between two subpopulations of isotopes: the marked population (usually the heavy isotope) and the unmarked population (usually the light population). Following the notations from López-Sepulcre et al. (2020), we define the state of those subpopulations in the network at time *t* by two vectors 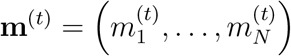 and 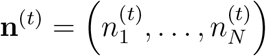 giving the marked and unmarked quantities of atoms, respectively, for each compartment. These vectors are related to the total pool sizes by **x**^(*t*)^ = **m**^(*t*)^ + **n**^(*t*)^ and the compartment labelled fractions are then defined as **z**^(*t*)^ = **m**^(*t*)^ ⊘ **x**^(*t*)^ where ⊘ is the element-wise division. In a typical tracer addition experiment, the observed time series are for **x**^(*t*)^ and **z**^(*t*)^ (not necessarily at the same time points), but for radioactive elements the observed time series can be simply **m**^(*t*)^.

The system of differential equations governing **m**^(*t*)^ and **n**^(*t*)^ is the same as for **x**^(*t*)^ above, the only differences being that **y**^(*t*)^ is split into **y**_**m**_^(*t*)^ and **y**_**n**_^(*t*)^ (the vectors defining the input of marked and unmarked matter, respectively, for each compartment) and that initial conditions usually differ for **m**^(*t*)^ and **n**^(*t*)^. Additionally, for radioactive tracers, the *λ* rate coefficients are adjusted to take into account the radioactive decay rate.

For a given set of parameter values, once the system of differential equations is solved and expected trajectories are calculated for **x**^(*t*)^ and **z**^(*t*)^, the observed time series for compartment sizes and labelled proportions are modelled as sampling and measurement errors around the expected trajectories. Several error distributions are implemented in `isotracer`. For example, for the observed size of compartment *i* at time *t*, one can assume a truncated Normal distribution with the mean being the expected compartment size and a coefficient of variation *cv* estimated by the model:

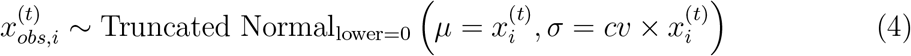

and for the observed labelled proportions one can assume, for example, a Beta distribution with mean the expected compartment labelled proportion and a precision parameter *ϕ* estimated by the model:

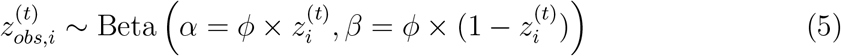

Other distributions can be a reasonable choice for the generation of observations from the expected trajectories. For example, Normal or Gamma distributions truncated at 0 can be a good approximation if their standard deviations are small and if the labelled proportions are well below 1, and are offered as an option in `isotracer`.

### 2.3 Modelling experimental design variations

The mathematical model described above can be adjusted to reflect specific features of a given tracer addition setup.

#### 2.3.1 The addition regime

A core component of modelling a tracer addition is the addition regime. The design of the addition regime is flexible, and additions can be done either at discrete time points (pulses, e.g. McRoy and Barsdate (1970); Rønnestad et al. (2000); Barthelemy et al. (2018)) or continuously over a given duration (continuous intervals or drips, e.g. de Goeij et al. (2013); Williams et al. (2016); Collins et al. (2016)). Additions can consist of only marked material, or a mix of marked and unmarked material. Finally, additions may be followed or not by an addition of fully unmarked material (chase, e.g. Simard et al. (1997); Carbone et al. (2007); Bacher et al. (2016)). All those regimes can be specified by an appropriate definition of the elements of vector 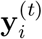.

Some source compartments might be considered to be in a steady state on the time scale of the experiment. This can be the case of dissolved nutrients in a stream which are constantly renewed by the water flow, or when the source compartment is extremely large relative to the amount of material which is transferred to consumer compartments (e.g. ants feeding on a large source of prepared medium). To model compartment *i* as being in a steady state, one needs to set all the coefficients of the corresponding *i*^*th*^ row of the **A** matrix to zero and *y*_*i*_(*t*) to zero: this will result in a constant value for 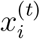 equal to the initial condition 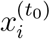.

#### 2.3.2 Overenrichment

Some sampled compartments can appear over-enriched compared to their source compartments. Over-enrichment happens when a receiving compartment appears more labelled than its source compartment. This occurs notably in ecosystem studies. Over-enrichment should not be possible according to the model, which assumes that the marked material is instantaneously mixed in the receiving compartment pool and that transfer from one compartment does not preferentially affect marked or unmarked material. However, some sampled compartments might actually represent the average of several sub-compartments, for example an active compartment involved in material flows and a refractory one which behaves as if isolated from the rest of the network on the time scale of the experiment. This is the case, for example, for a detrital compartment sampled on a stream bed, and which might represent a mix of algae and bacteria (the active portion), assimilating dissolved nutrients and preferentially grazed by invertebrate consumers, and a refractory portion of sediment and slowly decaying matter (e.g. wood and leaves), not involved in significant nutrient cycling in the span of the experiment. Invertebrate consumers can appear over-enriched compared to the average detrital compartment, but they are actually feeding mostly on the rich microbial fraction. For a split compartment *i*, comprised of an active and a refractory portions, modelling can be adjusted by adding a *π*_*i*_ parameter representing the active fraction of its pool at *t*_0_, and by only taking into account this active sub-pool when calculating material flows, but by taking into account the inactive, refractory portion when calculating expected total pool sizes and apparent labelled proportions (López-Sepulcre et al., 2020).

## 3 Package implementation

### 3.1 Overview

The model exposed above is implemented in isotracer using a Bayesian approach. The Stan program (Carpenter et al., 2017) and its R interace (the rstan package, Stan Development Team (2020)) are used for implementation of the MCMC. Stan uses a Hamiltonian Monte Carlo (HMC) algorithm to sample parameter posteriors with MCMC sampling. The HMC sampling allows for efficient and robust sampling of the posterior, and Stan output provides diagnostics to check that the sampled Markov chains behave appropriately. The Stan model first calculates the expected compartment trajectories for a set of parameter values (using the Euler method with constant step size *dt* to numerically solve the system of differential equations), then the likelihood is calculated by comparing observations to those expected trajectories. Note that, while the package user can choose the *dt* time step used during integration, no extensive check of numerical accuracy is performed during the numerical solving, and neither the ODE solvers nor the matrix exponential tools provided by Stan are used. This approach allows for greater speed of model evaluation and is expected to perform robustly when a correct *dt* time step and reasonable parameter priors are chosen, so that only a small fraction of the material from each compartment is transferred during a time step. In practice, a good rule of thumb is to try to keep the product between *dt* and the highest sampled values of any rate coefficient parameter below 0.1. In all cases post-run diagnostics should be performed by checking that the posterior predictions make sense and that re-running the model with smaller *dt* values does not change the posterior substantially.

### 3.2 Interface overview

The isotracer package separates the process of modelling tracer additions into two steps: (1) model definition, and (2) model fitting. This is due to the the amount of information required to define a network model, namely: network topology, initial conditions, observations, and priors. Figure 2 shows an overview of the typical workflow when using `isotracer` and Table 1 presents the main functions that are used in the model building step.

**Table 1:** `isotracer` functions used to define a network model (valid for version 1.0).

| Function | Role | Example | Notes |
| --- | --- | --- | --- |
| Definition of network properties |  |  |  |
| `new_networkModel()` | create a new, empty network model | `m <- new_networkModel()` | * |
| `set_topo()` | define a network topology | `m <- set_topo(m, "NH4 -> algae -> grazer")` | * |
| `set_init()` | define initial conditions | `m <- set_init(m, inits, comp = "compartment",`<br>`size = "biomass", prop = "fraction",`<br>`group_by = "stream_id")` | *a |
| `set_obs()` | define observations | `m <- set_obs(m, obs, time = "time_days")` | *a |
| `set_steady()` | set steady-state compartment | `m <- set_steady(m, "NH4")` |  |
| `set_split()` | set split compartment | `m <- set_split(m, "algae")` |  |
| `set_half_life()` | set a half-life (for radioactive tracer) | `m <- set_half_life(m, hl = 34.152)` |  |
| `add_pulse_event()` | set pulse and drip events | (see Trinidadian stream case study for examples) | b |
| Definition of statistical properties |  |  |  |
| `add_covariates()` | define categorical covariates | `m <- add_covariates(m,`<br>`upsilon_NH4_to_algae ~ stream_id)` |  |
| `set_size_family()` | set distribution for observed pool sizes | `m <- set_size_family(m, "normal_sd")` | c |
| `set_prop_family()` | set distribution for observed proportions | `m <- set_prop_family(m, "beta_phi")` | c |
| `set_prior()` | set prior distributions for parameters | `m <- set_prior(m, hcauchy(0.1),`<br>`"lambda_algae")` | d |

\* Required to build a runnable model.
a The example assumes that both `inits` and `obs` have columns named `compartment`, `biomass`, `fraction`, and `group_id`, and that `obs` also has a column named `time_days`.
b (Steady) drips can be defined by applying pulse events to steady state compartments.
c Default families are normal distributions for sizes and gamma distributions for proportions.
d Default priors are a half-Cauchy prior with a scale of 0.1 for all parameters except for active fractions of split compartments, for which the default is a uniform prior on $[0, 1]$ .

**Figure 2:**
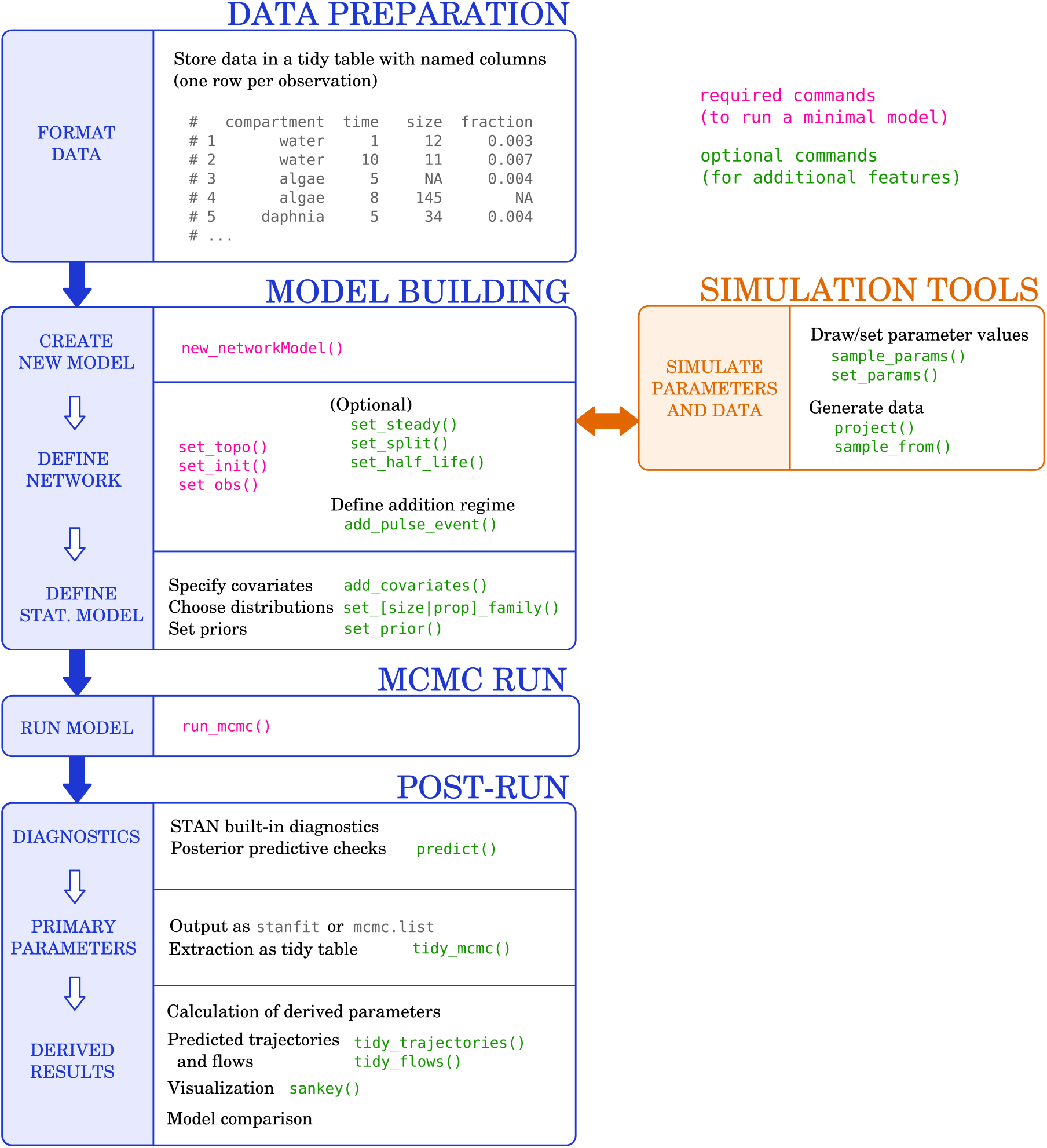
Workflow overview for network modelling with isotracer.

The package functions have been designed for convenient use with the pipe operator `%>%` provided by the `magrittr` package and familiar to `tidyverse` users, but can also be used without the pipe. A new, empty network model is initialized with the `new_networkModel()` function. This function returns a `tibble` data frame with zero rows and the supplementary class `networkModel`, ready to be populated as the user defines the model in subsequent steps. The `isotracer` package stores its user-facing objects using a tidy table approach (Wickham, 2014) as much as possible to make them easy to inspect. However, the user should use the package functions to modify a model to ensure it results in a valid model. Each row of a `networkModel` object corresponds to one network replicate. It is often necessary to be able to handle several replicates of a similar network system (e.g. modelling replicate sampling locations in a lake, modelling several transect locations along a stream, sampling several plants in a study of nutrient allocation, or modelling the removal of an injected drug in the bloodstream of several patients). All rows (i.e. all network replicates) share the same network topology, and each row can be considered either as a simple replication unit with its own initial conditions and observations, but sharing the parameter values that govern flows with all other replicates, or as a level of a treatment where categorical covariates are taken into account when estimating the model parameters, which can differ across treatments in this case. In other words, defining parameters as same or different among replicates defines whether there are different treatment or factor levels.

### 3.3 Model definition

#### 3.3.1 A minimum model

The usage of the functions that define a network model (Table 1) is presented in more details in the case studies and the corresponding appendix below. Here we give an overview of the minimum steps required to define a minimum executable model, and of the optional steps which allow fine-tuning model definition.

At least four functions have to be used to define a network model (the functions with an asterisk in the last column of Table 1):

- `new_networkModel()` to initialize a new, empty `networkModel` object.
- `set_topo()` to define the network topology of a `networkModel` (i.e. the network compartments and their connections).
- `set_init()` to provide the initial conditions (total size and labelled proportion) for each compartment. It takes a data frame with columns for compartment identity, size, and labelled proportion and accepts supplementary grouping variables used to define replicates.
- `set_obs()` to provide the observations (sampling time, total size, labelled proportion) for each compartment. It accepts a data frame in the same format as `set_init()`, with an extra column for time.

`isotracer` provides a default half-Cauchy prior with a scale of 0.1 for all parameters to be sampled (except for active fractions of split compartments, for which the default is a uniform prior on [0, 1]). Hence, a `networkModel` is technically already executable at this stage. However, the user should ensure that appropriate priors are used rather than relying blindly on the default ones, which might be inappropriate for the model at hand, depending on the time units used in the data provided by the user.

#### 3.3.2 Additional network properties

Depending on the network system that is being modelled, additional network properties can be set using the following functions:

- `set_steady()` to define which compartments should be considered at a steady state. The total size and labelled fraction of compartments at steady state do not change during the calculation of expected compartment trajectories (except if a pulse event is defined for them). This allows modelling, for example, compartments which are constantly replenished (such as dissolved nutrients in flowing stream water) or whose flows are insignificant relative to the time scale of the experiment.
- `set_split()` to define which compartments should be modelled as comprised of an active portion and a refractory portion.
- `set_half_life()` to define the half-life of radioactive isotopic tracers. This allows to take into account the radioactive decay of the tracer in the estimate of flows. By default, `isotracer` assumes that a stable tracer is used and no half-life is applied in the model.
- `add_pulse_event()` to add pulse events to specific network compartments. A pulse event is defined by an event time and by the quantities of labelled and unlabelled material added to the compartment at that time. In addition to allowing to define versatile “pulse” designs, this function can also be used to define “drip” designs when used along `set_steady()`: a pulse event applied to a steady-state compartment will result in a new steady state for this compartment, and “drip-on” and “drip-off” phases can thus be defined using a succession of pulse events to adjust the steady state of a compartment.

#### 3.3.3 Statistical model properties

All the functions above are used to define network properties. The statistical properties of a network model are defined with the following functions:

- `add_covariates()` to define categorical covariates with fixed effects on the estimated parameters.
- `set_size_family()` to define the distribution used to model measurement and sampling error when calculating the likelihood of observed compartment sizes compared to expected trajectories. Implemented distributions are:

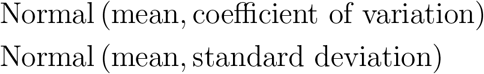
- `set_prop_family()` to define the distribution used to model measurement and sampling error when calculating the likelihood of observed labelled fractions compared to expected trajectories. Implemented distributions are:

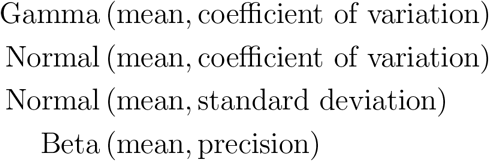
- `set_prior()` to set prior distributions for individual model parameters. Implemented priors are half-Cauchy, Normal, Uniform, and scaled Beta distribution. All priors are truncated to 0. Additionally, a “constant” prior is available to fix the value of some parameters during MCMC.

### 3.4 MCMC and posterior analysis

Once a network model is defined as explained above, MCMC sampling is performed with the `run_mcmc()` function. This function returns an object of class `mcmc.list` as implemented by the coda package, and which can be used directly by many R packages designed for Bayesian analyses (e.g. `bayesplot`). Optionally, `run_mcmc()` can be called with the `stanfit = TRUE` argument. In this case it will return the raw `stanfit` object produced by Stan. This is especially useful when Stan produces errors or warnings during the MCMC sampling: the `stanfit` object can be used for diagnostics (e.g. with the `shinystan` package) as explained in the Stan documentation (Carpenter et al., 2017). Once a model runs without any error or warning from Stan, the user is expected to use the default output from `run_mcmc()` (the `mcmc.list` object). Table 2 provides an overview of the main functions available for post-run processing of `mcmc.list` objects.

**Table 2:**
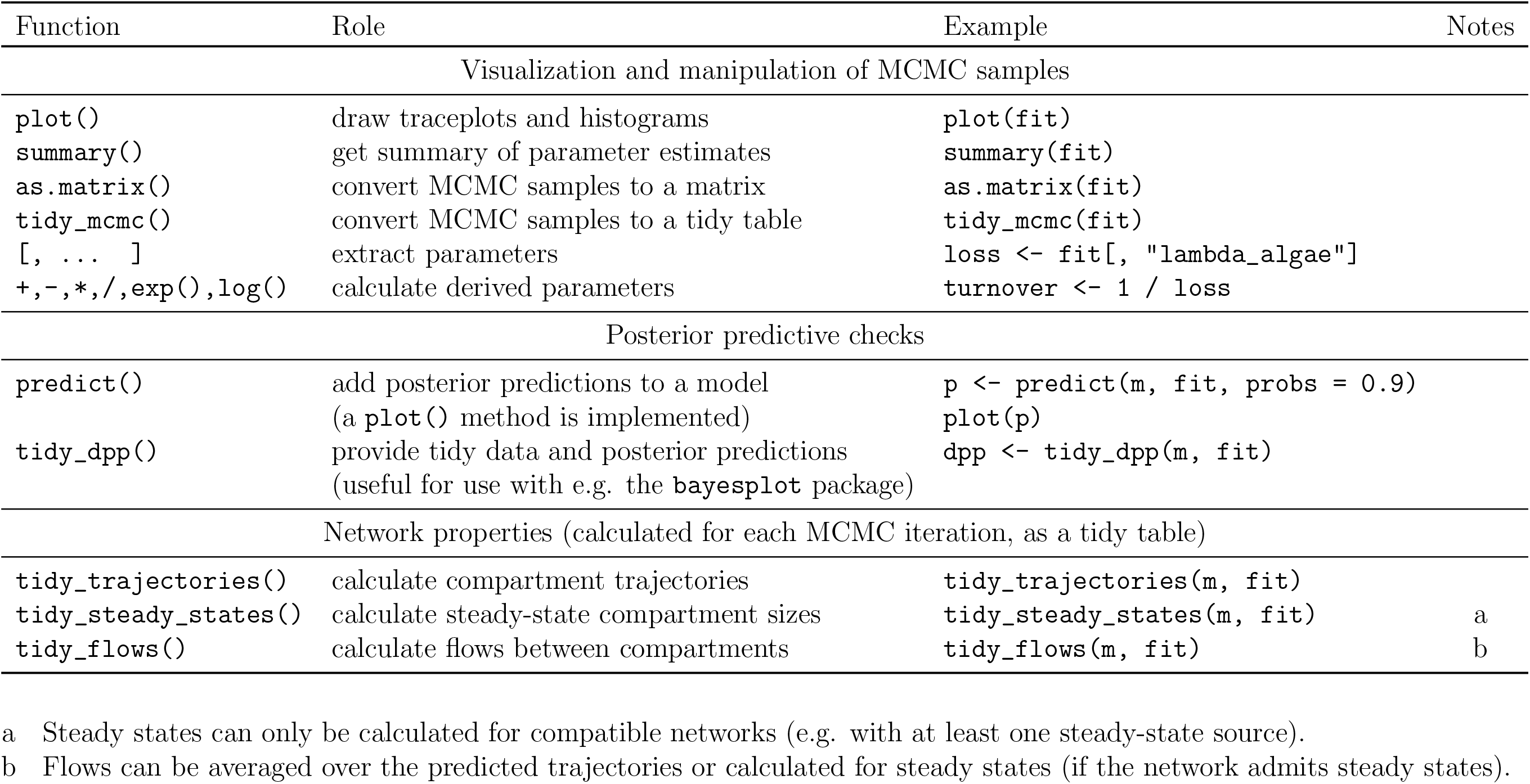
`isotracer` post-run functions typically used on the `mcmc.list` output of `run_mcmc()` (valid for version 1.0). The examples below assume that MCMC sampling was run on a network model m with for example: `fit <− run_mcmc(m)`.

| Function | Role | Example | Notes |
| --- | --- | --- | --- |
| Visualization and manipulation of MCMC samples |  |  |  |
| `plot()` | draw traceplots and histograms | `plot(fit)` |  |
| `summary()` | get summary of parameter estimates | `summary(fit)` |  |
| `as.matrix()` | convert MCMC samples to a matrix | `as.matrix(fit)` |  |
| `tidy_mcmc()` | convert MCMC samples to a tidy table | `tidy_mcmc(fit)` |  |
| `[, ... ]` | extract parameters | `loss <- fit[, "lambda_algae"]` |  |
| `+, -, *, / , exp(), log()` | calculate derived parameters | `turnover <- 1 / loss` |  |
| Posterior predictive checks |  |  |  |
| `predict()` | add posterior predictions to a model<br>(a `plot()` method is implemented) | `p <- predict(m, fit, probs = 0.9)`<br>`plot(p)` |  |
| `tidy_dpp()` | provide tidy data and posterior predictions<br>(useful for use with e.g. the `bayesplot` package) | `dpp <- tidy_dpp(m, fit)` |  |
| Network properties (calculated for each MCMC iteration, as a tidy table) |  |  |  |
| `tidy_trajectories()` | calculate compartment trajectories | `tidy_trajectories(m, fit)` |  |
| `tidy_steady_states()` | calculate steady-state compartment sizes | `tidy_steady_states(m, fit)` | a |
| `tidy_flows()` | calculate flows between compartments | `tidy_flows(m, fit)` | b |

a Steady states can only be calculated for compatible networks (e.g. with at least one steady-state source).
b Flows can be averaged over the predicted trajectories or calculated for steady states (if the network admits steady states).

The `isotracer` package provides additional methods for the `mcmc.list` class to facilitate the manipulation of such objects. For example, a plot method provides a compact way of visualizing MCMC traces and their histograms, and MCMC traces for derived parameters can be easily calculated from the primary parameters returned by `run_mcmc()` via methods implementing the common mathematical operators (+, −, ×, /, exp, log) for `mcmc.list` objects.

To assess model validity, `isotracer` provides functions to perform posterior predictive checks. A `predict()` method is provided for `networkModel` objects, which takes a model fit (the `mcmc.list` output from `run_mcmc()`) and calculates predicted trajectories based on the parameter posteriors. A `tidy_dpp()` funtion is provided for tidy data and posterior prediction calculation: its output contains the observed data as a y list element and the corresponding posterior predictions as a `y_rep` list element, which can be fed directly into the posterior predictive check functions provided by bayesplot, such as `ppc_dens_overlay()` or `ppc_intervals()`. The output of `tidy_dpp()` also contains a `vars` list element which can be used to provide the time points to the bayesplot plotting functions, and facetting variables when grouping factors were used in the model.

In addition to derived parameters and posterior predictions, `isotracer` can be used to calculate network properties in the particular case of a network admitting equilibrium states. A network can admit an equilibrium state if all *λ* parameters are set to 0 (no material is lost from the network system), or if at least one compartment is set to a steady state and not all *λ* are set to zero (a balance can be reached between network inflows and out-flows). Equilibrium states are calculated using eigenvectors of the transfer matrix. The calculation is performed by the `tidy_steady_states()` function, which will return the compartment size estimates at equilibrium for each MCMC iteration. Similarly, the `tidy_flows()` function can calculate the flows at equilibrium for each iteration, or the average flows over the experiment duration if the network does not admit an equilibrium state. More generally, the `tidy_trajectories()` function will return the full compartment trajectories for each MCMC iteration if they are needed for any downstream analysis.

Finally, since the package can predict compartment trajectories for a given set of parameter values, it can also be used to simulate experiments and generate datasets. The package provides functions to sample parameter values from their priors (`sample_params()`), to calculate trajectories for a set of parameter values (`project()`) and to generate predicted datasets including measurement and sampling error (`sample_from()`). Those functions can be used to optimize experimental designs before running costly and time-consuming real-life experiments, or to check parameter identifiability for a given design.

## 4 Case studies

The following case studies demonstrate the use of the `isotracer` package. In the first case study on *Arabidopsis* protein turnover, we introduce basic concepts such as the use of covariates, and the calculation of derived parameters. In the second case study on *Zostera* phosphate uptake, we demonstrate how to model radioactive isotopes, and how to compare derived parameters to a reference value. Finally, the study of the nitrogen cycle in a Trinidadian stream illustrates how to model drip experiments, non-homogeneous compartments, and how to perform model comparison. These case studies also illustrate the diversity of addition regimes that can be used in tracer additions (Figure 3). All datasets used in the case studies are available from the `isotracer` package.

**Figure 3:**
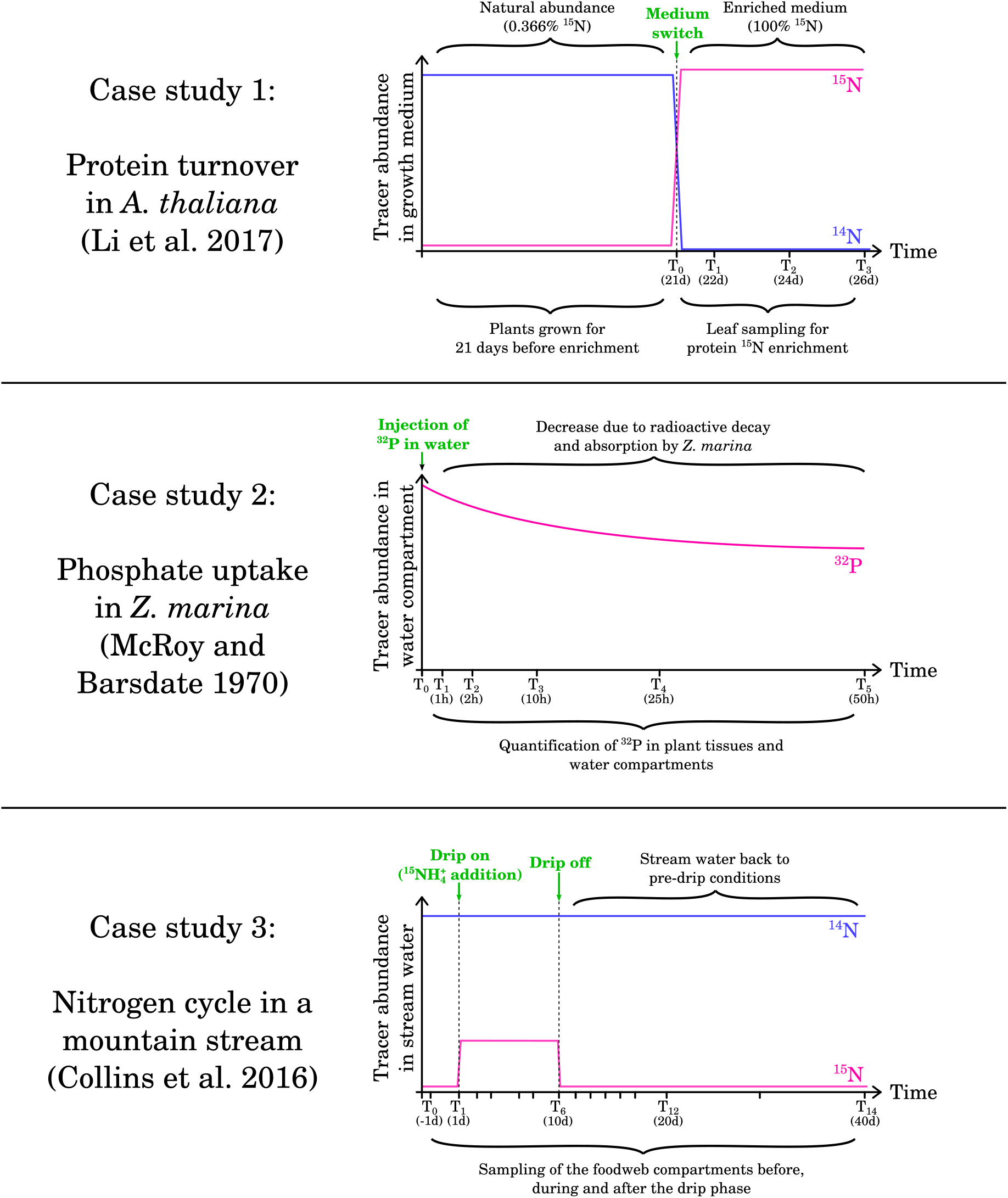
Overview of the addition regimes used in the three case studies.

### 4.1 Protein turnover in *Arabidopsis thaliana*

Well-regulated protein synthesis and degradation are crucial for organism development and maintenance. In a study by Li et al. (2017), ^15^N labelling was used to measure the degradation rate for 1228 proteins in *Arabidopsis thaliana* leaves from live plants. This experiment aimed at measuring the in vivo degradation rate of proteins in young plants (21-day old) in three different leaf tissues (3rd, 5th and 7th leaves). Obtaining turnover estimates for a large number of proteins allowed the authors to examine the determinants of protein degradation rates, such as protein domains and involvement in protein complexes, and to identify proteins for which degradation rates differed between leaves. Here, we demonstrate how to estimate turnover rates for a handful of proteins from the original study using `isotracer`, and how to compare those rates across leaf tissues.

In the original study, seeds were grown on a medium with naturally abundant N isotopes (mostly ^14^N) and switched to a ^15^N medium after 21 days. Leaves 3, 5, and 7 were taken from plants at 21 days (t0), 22 days (t1), 24 days (t2) and 26 days (t3) (Figure 3). Each sample consisted of four leaves (from four individual plants) pooled together into one biological replicate. For each leaf type and sampling time, three such replicates were collected. Leaf samples were used for protein separation on gel, digestion and mass spectrometry analysis in order to identify proteins and estimate their labelled fraction. In addition, a similar experiment was performed in which the seeds were grown only on natural N isotope medium (mostly ^14^N) and leaves were sampled at similar time points (without pooling multiple individuals). Those leaves were used in chromatography and mass spectrometry analysis to estimate relative abundance changes for individual proteins.

The data from Li et al. (2017) is available as a Dryad repository (Li et al., 2018) and the `isotracer` package ships several tables derived from this dataset. Here, we only focus on a handful of proteins and estimate their turnover in *Arabidopsis* leaves. The proteins we selected are RBCL (ribulose-bisphosphate carboxylase or “rubisco”, the most abundant protein in leaves in ppm), CPN60A (chaperone involved in rubisco folding), PSBA (protein of the photosystem II), THI1 (thiazole biosynthetic enzyme), and PGK1 (phosphoglycerate kinase 1). Several ways exists in which the biological question at hand can be answered with `isotracer`. For illustrative purposes, we choose to use a network model where the topology is extremely simple and each of the five protein compartments receives nitrogen “directly” from the growth medium (Figure 4A). This topology is replicated in three instances, one per leaf tissue, and leaf identity is used as a covariate in the model so that leaf-specific parameter values are estimated. This approach assumes that no exchange of nitrogen across proteins or across leaves occurs once nitrogen is incorporated into a protein. Finally, we assume that the N content in the medium is so large that it is constant compared to the protein pools, so we set it to a steady state in the model (Figure 4A). This is equivalent to the zero-order process assumed by Li et al. (2017) for protein synthesis.

**Figure 4:**
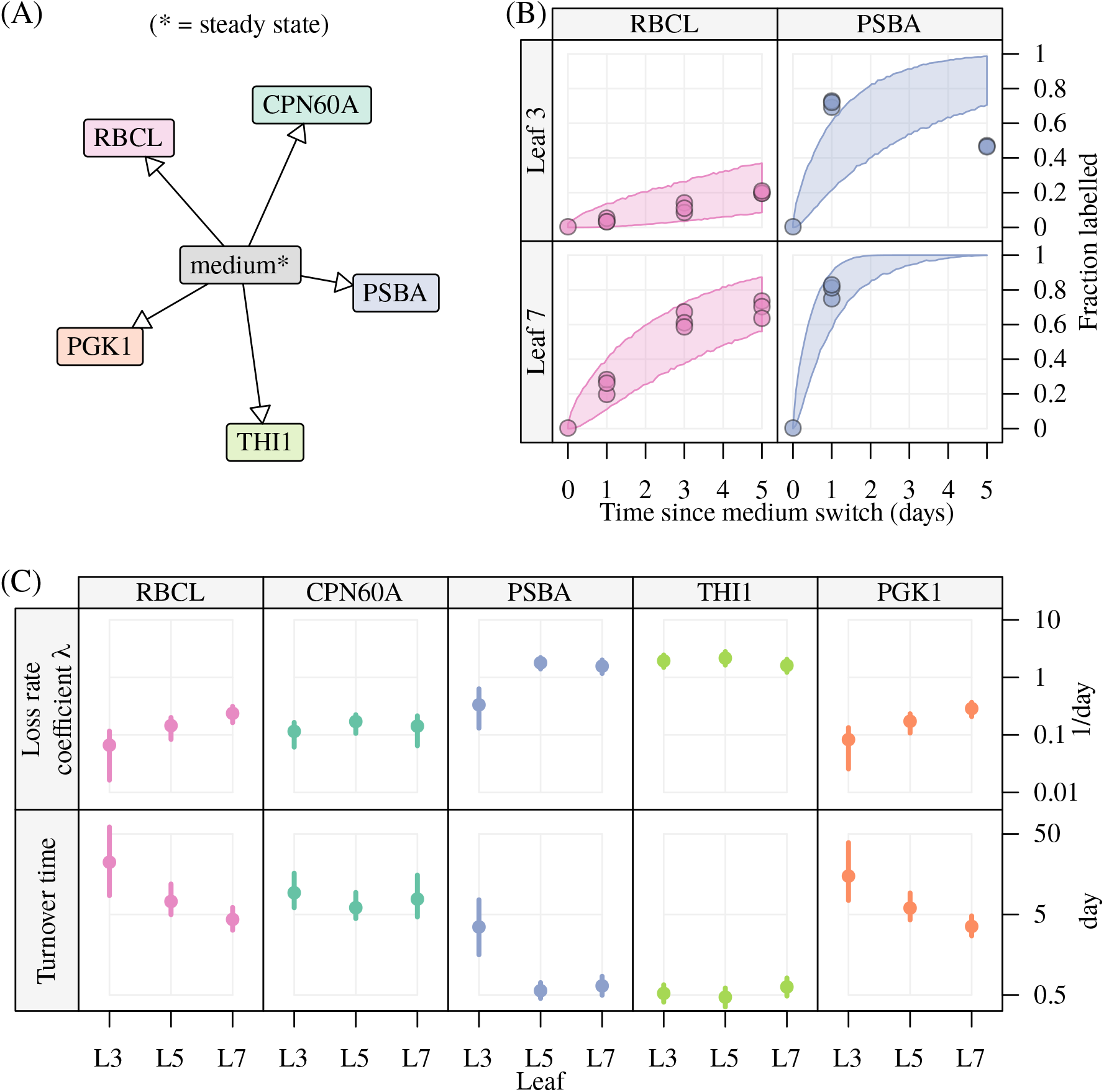
Modelling of protein degradation in *Arabidopsis thaliana* leaves based on Li et al. (2017). (A) Network topology used in our model. (B) Examples of posterior predictive checks for compartment enrichment (data points and 90% posterior prediction intervals). (C) Estimates for loss rate coefficients and turnover times for proteins across different leaves (posterior means and 95% credible intervals). Since the protein compartments do not transfer nitrogen to other compartments in the network model, their turnover times are simply the inverse of their *λ* parameters.

From a modelling perspective, the dataset from Li et al. (2017) contains all the data needed for an analysis with `isotracer`: a time series for compartments enrichment (labelled fractions) and a time series for compartments sizes (relative protein abundances). We model the network trajectories from the instant the medium was switched. We assume that the starting labelled fractions for all protein pools are the standard ^15^N abundance (0.3663%); for the medium, which is set to steady state in our model, we use a dummy relative abundance of 1 and a labelled fraction of 1 since almost all its N is ^15^N during the experiment. Once the MCMC sampling is complete, an important step is to check that the fitted model can correctly predict the original observations using a posterior predictive check (Figure 4B). A posterior predictive check consists in generating predicted data from the parameter posterior distribution, and comparing this predicted data with the actual observations used to fit the model. For example, if we use a 90% probability level, we would expect the predicted trajectory envelopes to contain about 90% of the original observations. Figure 4B shows this check for the labelled proportions of some compartments.

One of the aims of the original paper was to estimate the turnover rate of as many proteins as possible in the three leaf types, and compare those rates across proteins and leaves. Here, we estimated the turnover rates *λ* for five proteins, in three leaf types (since each protein pool is not transferring nitrogen to any other compartment, its turnover rate is equal to *λ*) (Figure 4C). A powerful aspect of Bayesian MCMC is that we can calculate posterior distributions for derived parameters by doing calculations on the primary parameters sampled during MCMC. For example, turnover times are the inverse of turnover rates, and Figure 4C illustrates posteriors for turnover time which were derived from the turnover rates.

### 4.2 Phosphate uptake in *Zostera marina*

Eelgrass can make dense populations in shallow water areas and have an important role in nutrient cycles in the ecosystems they inhabit. McRoy and Barsdate (1970) used controlled laboratory conditions to characterize the uptake of phosphorus by eelgrass plants (*Zostera marina*) from the surrounding water or sediment and its transfer in the plant tissues under light and dark conditions. Their experimental setup consisted in individual eelgrass plants kept in closed jars, with a watertight septum separating the upper and the lower compartments. The upper compartment contained the leaves and stems of the plant, while the lower compartment contained the roots and rhizome. ^32^P was added as phosphate to either the upper or lower water compartment at the beginning of the experiment, and jars were kept in either light or dark conditions. At several time points over the course of two days, individual plants were taken from the experiment and ^32^P abundance in the water and in the plant tissues was quantified by measuring radioactive decay (cpm/mg of dry material) (Figure 3). Here, we estimate the phosphate flows in a simplified network with four compartments: sediment water (“lower water”), free water (“upper water”), roots + rhizome (“lower plant”), and leaves + stem (“upper plant”). To make the model more amenable, we neglect the release of phosphorus by the plant into the water and only consider four connections in the network model: from upper water to leaves and stem, from lower water to roots and rhizome, and bidirectional flow between leaves and stem (upper plant) and roots and rhizome (lower plant). We examine the light effect by using the light treatment as a covariate in our model, while the addition of ^32^P to either the upper or the lower compartment is considered as a replication within light treatment. In other words, we assume that the rate coefficients defining the phosphorus flows do not depend on the compartment where ^32^P was added, but that the light conditions could have an effect on the rate coefficients. The radioactive decay of ^32^P is automatically taken into account by `isotracer` by adding the appropriate decay rate to the estimated loss rate coefficient of each compartment once the half-life has been specified in the model. Note that since only ^32^P (not total P) was measured in these experiments, our model considers only ^32^P pools and we fixed the labelled fractions of all observations to 1 (fully labelled).

We compiled the data from McRoy and Barsdate (1970) from the original article figures and tables. We used some approximations to convert the reported values to usable time series. Notably, the reported data for each time point was in cpm/mg of dry weight, and we used a single mean value of each tissue total dry weight per treatment to convert those values into ^32^P quantities per compartment for the purpose of this illustrative case study. Given those approximations, the results reported here should be taken as an illustration of the use of `isotracer`, rather than as a biologically precise report on eelgrass physiology.

After fitting the model to determine the transfer rate parameters, we estimated the average flows between compartments over the time of the experiment, in each light treatment (Figure 5A). The Sankey plot shown in this figure is a useful way to visualize flows across a network, but it should be noted that it only includes point estimates of the flows. For a statistically robust comparison between treatments, we can perform calculations on the MCMC values of the primary parameters to build posteriors for derived parameters. For example, to test if the uptake rate coefficients differed between light treatments, we calculated the posteriors for their ratios between light and dark conditions (Figure 5B). Based on Figure 5B, only the coefficient for phosphate uptake from upper water to the leaves and stem increased significantly in light. For the other coefficients, the 95% credible intervals were so large that they overlapped with a ratio value of 1. This large uncertainty is due to the variability of the original data and the blunt assumptions that we made when we converted the original data to usable time series. We expect that a dataset where total tissue dry weight were estimated at the same time as cpm/mg dry weight would allow to reduce the uncertainty a lot. However, this illustrates the ability of our method to estimate uncertainty in derived parameters and to test rigorously for differences between treatments.

**Figure 5:**
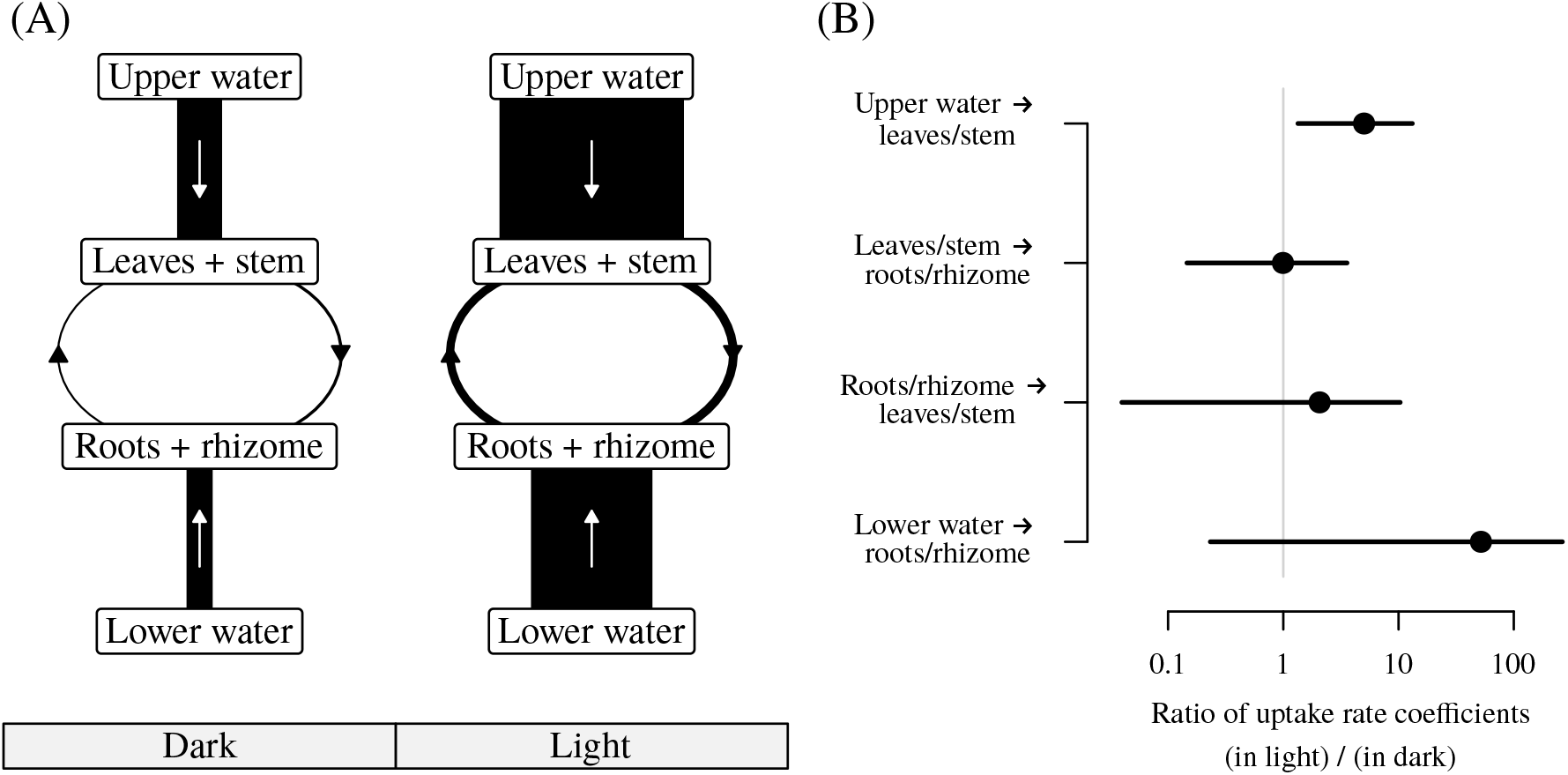
Modelling of phosphate absorption in *Zostera marina* tissues under dark and light conditions using data from McRoy and Barsdate (1970). (A) Sankey plot summarizing estimated average phosphate flows between study system compartments. The width of the connectors is proportional to the estimated flows. In the original experiment, upper and lower water compartments are separated by a septum. (B) Mean estimates and 95% credible intervals for the ratios of the uptake rate coefficient values between light and dark conditions.

### 4.3 Nitrogen cycle in a Trinidadian mountain stream

Tracer addition experiments have been used profitably in ecology to identify the strength of trophic connections and to quantify nutrient cycling in ecosystems. In a study by Collins et al. (2016), the authors quantified the nitrogen exchanges in the foodwebs of mountain streams in Trinidad, from the dissolved nutrients (NH_4_^+^ and NO_3_^−^) to the invertebrate consumers. They examined the effect of light conditions and of the presence/absence of a fish consumer on the nitrogen dynamics in the foodweb. Data was collected by dripping ^15^N-enriched ammonium into two streams in Trinidad, and samples from each foodweb compartment were taken during the drip and after the drip in several transects in each stream. The transects were located at different locations downstream of each drip. The drip phase lasted 10 days, and the post-drip phase lasted 30 days (Figure 3). Compartment enrichment was measured by determining ^15^N/^14^N ratios by mass spectrometry, while compartment sizes were estimated in mgN/m^2^ using various field sampling techniques depending on the compartment type (dissolved, primary producer, invertebrate). Streams differed in their exposure to sunlight (natural versus trimmed canopy) and transects within streams differed in the presence/absence of guppies (*Poecilia reticulata*). The dataset used in Collins et al. (2016) is shipped with `isotracer` as the `lalaja` table. In this case study, we will use the data from only one stream and a subset of the original foodweb for simplicity, but a full analysis including both streams and using the framework used in `isotracer` was presented in López-Sepulcre et al. (2020). We show how `isotracer` can be used to test for the existence of a candidate trophic link by comparing models, to determine active and refractory portions of inhomogeneous compartments, and to estimate the relative proportions of inputs into a given compartment.

We ran two models using the topology presented in Figure 6A. Those models differed only by the inclusion or not of a candidate trophic link between the grazer genus *Psephenus* and the predator genus *Argia* (dashed arrow in Figure 6A). Dissolved nutrients were set to a steady state given that they were renewed by the stream flow, and we allowed the two primary producer compartments, the algal cover on rock subtrate (epilithon) and the fine benthic organic matter (FBOM), to be modelled as split (inhomogeneous) compartments given that the field sampling cannot distinguish between active and refractory portions of those compartments. We used data from three transects from one stream (Upper LaLaja, for which the canopy cover was thinned). The transects differed in their effective addition regime due to their different distances from the drip source, but otherwise were used as simple replicates in our model (i.e. they shared the same parameter values). We used DIC to compare the models: the ΔDIC between the models was relatively small (2.73), and suggested that the model without the candidate trophic link fitted the data better, with a DIC-derived weight of 80% for this model (López-Sepulcre et al., 2020). Posterior predictive checks were statisfactory (Figure 6B).

**Figure 6:**
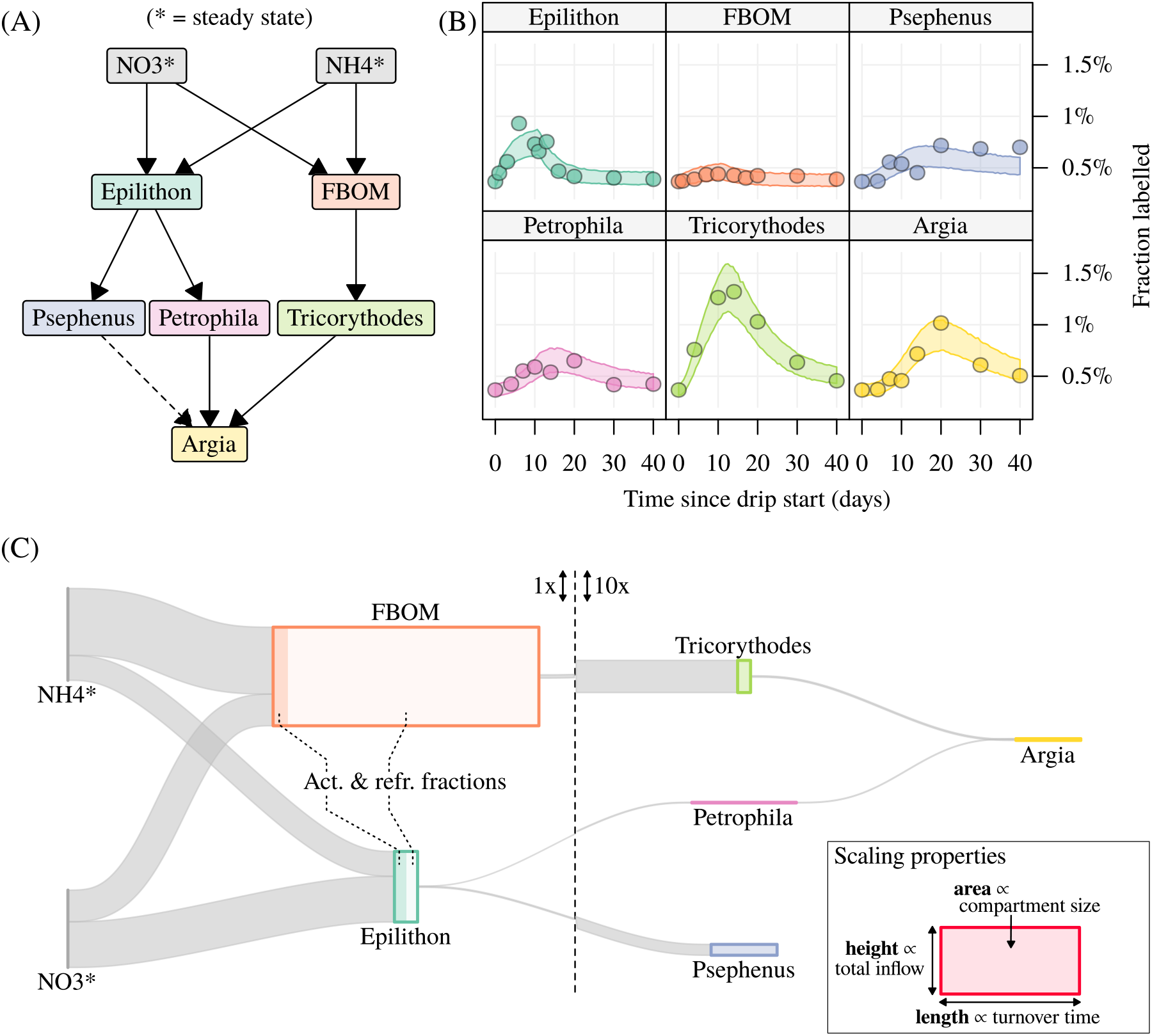
Modelling on nitrogen transport across foodwebs in Trinidadian mountain streams using data from Collins et al. (2016). (A) Foodweb topology used in our case study. The dashed arrow depicts the candidate trophic link that is tested by model comparison. (B) Examples of posterior predictive checks for the labelled fraction data in the first transect of the stream (data points and 90% posterior prediction intervals). (C) Sankey plot showing the estimated nitrogen flows across compartments in the first transect. Connecting ribbon widths are proportional to estimated flows. For epilithon and FBOM, an active portion of the compartment was estimated by our model. Vertical scale is expanded in the right half of the plot for readability.

As for the *Zostera* case study, we can use a Sankey plot to visualize flows between compartments (Figure 6C). In this case, the area of a given compartment is proportional to the compartment size (a quantity of nitrogen) while the thickness of the connecting ribbons is proportional to the flows (a quantity of nitrogen per time unit). It results from this that the length of a given compartment is proportional to its turnover time (per time unit). The Sankey plot clearly shows the difference in flows between nutrients and primary producers, compared to between primary producers and upper trophic levels. The estimated active portions in epiliton and FBOM were 0.51 (95%CI 0.25-0.77) and 0.05 (95%CI 0.01-0.14), respectively. Being able to estimate an active portion instead of having to set it to a fixed value a priori is important when no prior knowledge is available to make a robust, educated guess about this value. It is also the case for proportions of different inputs: the modelling approach implemented in `isotracer` estimates proportions of different inputs into a compartment from the tracer data, rather than having to rely on estimates from e.g. gut content analysis or other independent studies. For epilithon, the proportion of nitrogen assimilated from NH_4_^+^ was 0.36 (95%CI 0.26-0.49) and from NO_3_^−^ 0.64 (95%CI 0.51-0.74). The high part of NO_3_^−^ is explained by the open canopy of this stream, while estimates in the other, closed canopy stream showed that epilithon favoured NH_4_^+^ when light was limiting (López-Sepulcre et al., 2020).

## 5 Concluding remarks: current limitations, extensibility, and future directions

The `isotracer` package implements a Bayesian approach to model tracer addition experiments and to estimate parameter uncertainty rigorously. Its primary aim is to allow researchers whose typical expertise might not cover statistical modelling of systems governed by ODE to perform rigorous and reliable statistical analyses, so that the valuable data produced by tracer addition experiments can be fully taken advantage of. The package allows for modelling of a variety of addition designs (pulse, drip, pulse-chase), can take into account both stable and decaying (e.g. radioactive) tracers, and can be used for model comparison to test hypotheses relating to network structure. The toolkit provided for post-MCMC analyses is versatile and can be used to calculate network-wide properties (and their uncertainties) such as flow estimates or compartments steady state sizes when applicable.

From a technical point of view, the numerical solver currently used is reasonably fast (running 2000 MCMC iterations for the Trinidadian stream case study presented here took 90 seconds on a 3.6 GHz processor with one core per chain). This performance is at the expense of automatic numerical accuracy checks and the solver must thus be used with some caution. Stan diagnostics can help detect situations where this approach fails in obvious ways, but it is strongly recommended in all cases to perform some extraneous MCMC runs with a few smaller time step values once the model behaves appropriately in order to check for numerical stability. Future versions could implement an option to use Stan’s ODE solver or matrix exponentials for increased robustness, at the cost of speed of execution, to facilitate this checking step.

From a modelling perspective, a future improvement could include the implementation of continuous covariates in addition to discrete ones. This would allow the study of phenomena such as size-specific uptake in consumers, or temperature-specific resource allocation. Isotope fractionation could also be taken into account: the current version of `isotracer` ignores it and assumes that the tracer and the corresponding “unmarked” material are governed by identical transfer rate coefficients. For ranges of isotope enrichment usually observed in trophic networks (e.g. *δ*^15^N typically increasing by about 3.4 ‰ per trophic level (Cabana and Rasmussen, 1994), but with large variations depending on the study system, e.g. Mill et al. (2007)), it is likely that this is a reasonable approximation given the very large enrichment caused by the experimental addition (e.g. *δ*^15^*N* up to about 2500 ‰ in some primary consumers in Collins et al. (2016), but as low as 40 ‰ in higher trophic levels further away from the drip source). However, to accurately take into account isotopic fractionation at higher trophic levels or due to other processes (Robinson, 2001), an option to model it when it is known to occur could be implemented.

Importantly, the current version of `isotracer` only models first-order reactions: the quantity of transported material per unit of time is calculated as the product of a constant rate coefficient and of the source compartment size. Such first-order reactions result in simple systems of differential equations, and offer in most cases a good approximation to the modelled network dynamics on the time scale of the experiment. However, in some scenarios, second-order reactions, or more complex equations, govern the material flows. In other cases, the rate coefficients are not constant over the time scale of the experiment (e.g. time-dependent rate of glucose uptake from blood-stream after injection). The `isotracer` package can be extended with more complex models to be able to model such cases, including cases where several sources must react together following stoichiometric constraints (e.g. kinetic flux profiling of known metabolic pathways, Yuan et al. (2006)). Such a task would involve extending both the Stan code (to calculate the expected compartment trajectories in the Stan model) and the R code (to provide the interface to those new models and to define the post-run methods used e.g. to perform posterior predictive checks on the Stan output). We hope that feedback from users interested in package extension with new models will help prioritize which models should be implemented in later versions.

In summary, we have presented a package that implements and extends López-Sepulcre et al. (2020)’s method to analyze isotope tracer addition experiments in a diversity of systems and levels of organization, and under a variety of experimental designs. This method allows for the quantification of uncertainty in the estimation of matter and nutrient flows, turnover times, and other derived properties. As such, it allows for rigorous statistical inferences in comparative or experimental contexts. The package in its current format already fulfills the needs of a majority of short-term experiments at typically studied scales, and it could be extended to accommodate new analyses and scenarios in future versions. We believe it represents a valuable tool to advance the study of the dynamics of chemical elements in tissues, organisms, and ecosystems.

## Supporting information

Appendix (tutorials for case studies)

## 6 Acknowledgments

We would like to thank Sarah Collins, Sebastiano De Bona, Rana El-Sabaawi, Swanne Gordon, Sophia Lambert, Alex Lee, and Steven Thomas for discussions. Sarah Collins and Jonathan Benstead helped test the package interface. Funding was provided by the Academy of Finland (grant number 295941).

## 7 Package availability

`isotracer` is available from https://matthieu-bruneaux.gitlab.io/isotracer/ and will be submitted to CRAN in the near future. The source code is available as a Git repository from https://gitlab.com/matthieu-bruneaux/isotracer.

