## Appendix (tutorials for case studies) for "isotracer: An R package for the analysis of tracer addition experiments"

### Appendix: Detailed Tutorials for the Presented Case Studies

2021-08-09

#### Contents

**Case study 1: Protein turnover in *Arabidopsis* (Li et al. 2017)** **2**

**Case study 2: Phosphorus uptake in eelgrass (McRoy and Barsdate 1970)** **12**

**Case study 3: Trinidadian streams (Collins et al. 2016)** **20**

**Computing environment** **33**

*Note: The tutorials presented here are also available in the* `isotracer` *package documentation.*

### Case study 1: Protein turnover in *Arabidopsis* (Li et al. 2017)

This case study estimates protein turnover in *Arabidopsis thaliana* from experiment where seeds are grown on $^{15}$N medium. The data is taken from the following article:

- Li, Lei, Clark J. Nelson, Josua Trösch, Ian Castleden, Shaobai Huang, and A. Harvey Millar. "Protein Degradation Rate in Arabidopsis Thaliana Leaf Growth and Development." The Plant Cell 29, no. 2 (February 1, 2017): 207–28. https://doi.org/10.1105/tpc.16.00768.

In this article, the authors use chromatography, gel electrophoresis and mass spectrometry to estimate the degradation rate of 1228 proteins in *Arabidopsis* leaves. The data from their study was deposited on Dryad:

- Li, Lei, Clark J. Nelson, Josua Troesch, Ian Castleden, Shaobai Huang, and A. Harvey Millar. "Data from: Protein Degradation Rate in Arabidopsis Thaliana Leaf Growth and Development." Dryad, 2018. https://doi.org/10.5061/DRYAD.Q3H85.

A subset of the Dryad dataset is included in the datasets shipped with `isotracer`.

#### Experiment and data

The experiment in Li et al. (2017) aims at measuring the in vivo degradation rate of proteins in young plants (21-day old) in three different leaf tissues (3rd, 5th and 7th leaves). Obtaining turnover estimates for a very large number of proteins enabled them to examine the determinants of protein degradation rates, such as protein domains and membership in protein complexes, and to analyze proteins for which degradation rates differ between leaves.

In brief, seeds were grown on a medium with naturally abundant N isotopes (mostly $^{14}N) and switched to a 15N medium after 21 days. Leaves 3, 5, and 7 were taken from plants at 21 days (t0), 22 days (t1), 24 days (t2) and 26 days (t3). Each sample consisted of four leaves (from four individual plants) pooled together into one biological replicate. For each leaf type and sampling time, three such replicates were collected.

Leaf samples were used for protein separation on gel, digestion and mass spectrometry analysis in order to identify proteins and estimate their labelled fraction.

In addition, a similar experiment was performed in which the seeds were grown only on natural N isotope medium (mostly 14N) and leaves were sampled at similar time points (without pooling multiple individuals). Those leaves were used in chromatography and mass spectrometry analysis to estimate relative abundance fold changes for individual proteins.

The data shipped with `isotracer` is:

- `li2017`: a table providing relative abundance and labelled fraction data for individual proteins, for different days and different leaf types
- `li2017_prots`: a table mapping the protein identifiers from `li2017` to their description
- `li2017_counts`: a summary table giving the number of relative abundance data points and of labelled fraction data points for each protein.

This is how the relative abundance and labelled fraction data looks like:

```
li2017
```

```
## # A tibble: 59,048 x 6
##    prot_id                   sample rel_abundance labeled_fraction time_day leaf_id
##    <chr>                     <chr>          <dbl>            <dbl>    <dbl> <chr>
##  1 AT1G01050.1               T0L3R3         1.29                NA        0 leaf_3
##  2 AT1G01050.1               T0L5R3         1.29                NA        0 leaf_5
##  3 AT1G01080.1,AT1G01080.2 T0L3R1         0.972               NA        0 leaf_3
##  4 AT1G01080.1,AT1G01080.2 T0L3R2         1.10                NA        0 leaf_3
##  5 AT1G01080.1,AT1G01080.2 T0L3R3         0.854               NA        0 leaf_3
##  6 AT1G01080.1,AT1G01080.2 T0L5R1         1.37                NA        0 leaf_5
##  7 AT1G01080.1,AT1G01080.2 T0L5R2         1.07                NA        0 leaf_5
```

```
##  8 AT1G01080.1,AT1G01080.2 T0L5R3          0.747               NA         0 leaf_5
##  9 AT1G01080.1,AT1G01080.2 T0L7R1          0.332               NA         0 leaf_7
## 10 AT1G01080.1,AT1G01080.2 T0L7R2          1.09                NA         0 leaf_7
## # ... with 59,038 more rows
```

In this case study, we will only focus on a handful of proteins and use `isotracer` to estimate their turnover in *Arabidopsis* leaves. These are the proteins we selected for this case study:

- ATCG00490.1 (**RBCL**, ribulose-bisphosphate carboxylase or “rubisco”, the most abundant protein in leaves in ppm)
- AT5G54770.1 (**THI1**, thiazole biosynthetic enzyme)
- AT2G28000.1 (**CPN60A**, a chaperone involved in rubisco folding)
- ATCG00020.1 (**PSBA**, a protein of the photosystem II)
- AT3G12780.1 (**PGK1**, phospohoglycerate kinase 1)

#### Building the model

##### Network topology

For a given protein, the network model describing nitrogen fluxes has three compartments: the N source the growth medium and the protein pool in the three leaves (3rd, 5th and 7th leaves). We assume that the proteins in each leaf incorporate nitrogen “directly” from the growth medium, and that no exchange of nitrogen across leaf pool occurs once nitrogen is incorporated in a protein.

We initialize the network model with the corresponding topology, which is very simple: each protein pool is incorporating N as the corresponding protein is synthetized.

```
m <- new_networkModel() %>%
  set_topo("medium -> RBCL, THI1, CPN60A, PSBA, PGK1")
```

We assumes that the N content in the medium is so large it is constant compared to the leaf pools and set it to a steady state in the model. This is equivalent to the zero-order process assumed by Li et al. for protein synthesis.

```
m <- m %>% set_steady("medium")
```

Finally, the protein labelling in this experiment can be very high (proteins with a rapid turnover can be 100% labelled after a few days). This means that the default gamma distribution used to model observed labelled fractions is not appropriate, and we use a beta distribution instead:

```
m <- m %>% set_prop_family("beta_phi")
```

With this family, the model parameter $\eta$ represents the precision parameter $\phi$ of a beta distribution.

Let’s have a look at our network topology:

```
ggtopo(m, layout = "stress")
```

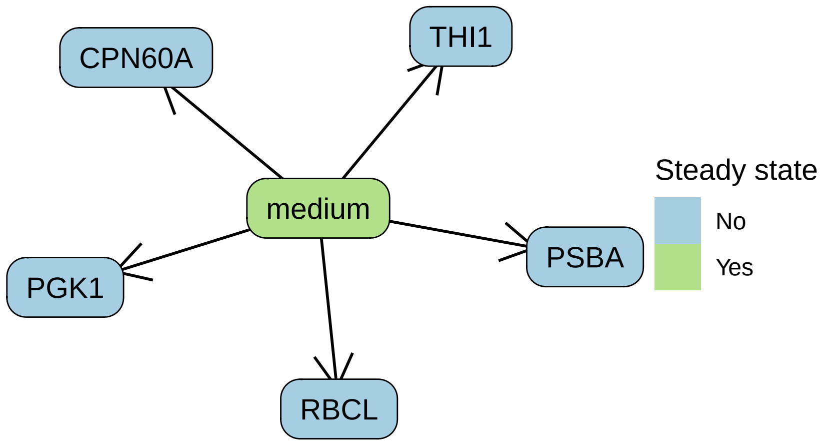


##### Initial conditions and observations

Let's get the data for those proteins. At the same time, we map the protein identifiers to something slightly easier to read, ad we also rename the protein ID column to `compartment` because we will have to add the medium nitrogen pool later:

```
prots <- c("ATCG00490.1" = "RBCL", "AT5G54770.1" = "THI1", "AT2G28000.1" = "CPN60A",
           "ATCG00020.1" = "PSBA", "AT3G12780.1" = "PGK1")
data <- li2017 %>%
  filter(prot_id %in% names(prots)) %>%
  mutate(prot = prots[prot_id]) %>%
  rename(compartment = prot) %>%
  select(compartment, leaf_id, time_day, rel_abundance, labeled_fraction)
data
```

```
## # A tibble: 284 x 5
##    compartment leaf_id time_day rel_abundance labeled_fraction
##    <chr>       <chr>      <dbl>         <dbl>            <dbl>
##  1 CPN60A      leaf_3         0         0.837               NA
##  2 CPN60A      leaf_3         0         0.818               NA
##  3 CPN60A      leaf_3         0         0.953               NA
##  4 CPN60A      leaf_5         0         1.40                NA
##  5 CPN60A      leaf_5         0         1.31                NA
##  6 CPN60A      leaf_5         0         1.73                NA
##  7 CPN60A      leaf_7         0         1.44                NA
##  8 CPN60A      leaf_7         0         2.03                NA
##  9 CPN60A      leaf_7         0         2.06                NA
## 10 CPN60A      leaf_3         1         0.774               NA
## # ... with 274 more rows
```

We get the initial conditions. Because there are several replicate, we will take their average as the starting condition:

```
inits <- data %>%
  filter(time_day == 0) %>%
  group_by(compartment, leaf_id, time_day) %>%
  summarize(rel_abundance = mean(rel_abundance), .groups = "drop")
inits
```

```
## # A tibble: 15 x 4
##    compartment leaf_id time_day rel_abundance
```

```
##    <chr>       <chr>      <dbl>         <dbl>
##  1 CPN60A      leaf_3         0         0.870
##  2 CPN60A      leaf_5         0         1.48
##  3 CPN60A      leaf_7         0         1.84
##  4 PGK1        leaf_3         0         1.12
##  5 PGK1        leaf_5         0         1.13
##  6 PGK1        leaf_7         0         0.785
##  7 PSBA        leaf_3         0         1.46
##  8 PSBA        leaf_5         0         1.36
##  9 PSBA        leaf_7         0         0.993
## 10 RBCL        leaf_3         0         0.993
## 11 RBCL        leaf_5         0         1.23
## 12 RBCL        leaf_7         0         0.944
## 13 THI1        leaf_3         0         0.878
## 14 THI1        leaf_5         0         1.23
## 15 THI1        leaf_7         0         1.24
```

Those initial conditions lack two things: the initial labelled fraction and the data for the medium nitrogen pool. We will assume that the starting labelled fraction is the standard 15N abundance (0.3663%). For the medium, which is in steady state, we use a dummy relative abundance of 1 and a labeled fraction of 1 because almost all the N is 15N during the experiment.

```
inits <- inits %>%
  mutate(labeled_fraction = 0.003663)
# We use the same medium specifications for all proteins
inits_medium <- tibble(compartment = "medium", rel_abundance = 1, labeled_fraction = 1) %>%
  crossing(inits %>% select(leaf_id) %>% unique())
# Put it all together
inits <- bind_rows(inits, inits_medium)
```

Now we can set those initial conditions in the model:

```
m <- m %>% set_init(inits, comp = "compartment", size = "rel_abundance",
                    prop = "labeled_fraction", group_by = "leaf_id")
```

```
# If we had the PaxDB abundance data we can convert the relative flows to
# absolute flows, in addition to having the turnover estimates.

# Another way to see things would be to have only two compartments, medium and
# protein, and use leaf and prot as grouping facto (probably not the best
# option).
```

We add the observations to the model:

```
m <- m %>% set_obs(data, time = "time_day")
```

Finally, we specify that all the rates will be dependent on the protein identity:

```
m <- m %>% add_covariates(upsilon + lambda ~ leaf_id)
```

This results in quite a few parameters to be estimated by the model:

```
params(m)
```

```
##  [1] "eta"                          "lambda_CPN60A|leaf_3"
##  [3] "lambda_CPN60A|leaf_5"         "lambda_CPN60A|leaf_7"
##  [5] "lambda_medium|leaf_3"         "lambda_medium|leaf_5"
##  [7] "lambda_medium|leaf_7"         "lambda_PGK1|leaf_3"
##  [9] "lambda_PGK1|leaf_5"           "lambda_PGK1|leaf_7"
## [11] "lambda_PSBA|leaf_3"           "lambda_PSBA|leaf_5"
## [13] "lambda_PSBA|leaf_7"           "lambda_RBCL|leaf_3"
## [15] "lambda_RBCL|leaf_5"           "lambda_RBCL|leaf_7"
## [17] "lambda_THI1|leaf_3"           "lambda_THI1|leaf_5"
```

```
## [19] "lambda_THI1|leaf_7"              "upsilon_medium_to_CPN60A|leaf_3"
## [21] "upsilon_medium_to_CPN60A|leaf_5" "upsilon_medium_to_CPN60A|leaf_7"
## [23] "upsilon_medium_to_PGK1|leaf_3"   "upsilon_medium_to_PGK1|leaf_5"
## [25] "upsilon_medium_to_PGK1|leaf_7"   "upsilon_medium_to_PSBA|leaf_3"
## [27] "upsilon_medium_to_PSBA|leaf_5"   "upsilon_medium_to_PSBA|leaf_7"
## [29] "upsilon_medium_to_RBCL|leaf_3"   "upsilon_medium_to_RBCL|leaf_5"
## [31] "upsilon_medium_to_RBCL|leaf_7"   "upsilon_medium_to_THI1|leaf_3"
## [33] "upsilon_medium_to_THI1|leaf_5"   "upsilon_medium_to_THI1|leaf_7"
## [35] "zeta"
```

#### Running the model

The default priors are half-Cauchy distributions with a scale of 1:

```
priors(m)
```

```
## # A tibble: 35 x 2
##    in_model             prior
##    <chr>                <list>
##  1 eta                  <hcauchy (scale=0.1)>
##  2 lambda_CPN60A|leaf_3 <hcauchy (scale=0.1)>
##  3 lambda_CPN60A|leaf_5 <hcauchy (scale=0.1)>
##  4 lambda_CPN60A|leaf_7 <hcauchy (scale=0.1)>
##  5 lambda_medium|leaf_3 <hcauchy (scale=0.1)>
##  6 lambda_medium|leaf_5 <hcauchy (scale=0.1)>
##  7 lambda_medium|leaf_7 <hcauchy (scale=0.1)>
##  8 lambda_PGK1|leaf_3   <hcauchy (scale=0.1)>
##  9 lambda_PGK1|leaf_5   <hcauchy (scale=0.1)>
## 10 lambda_PGK1|leaf_7   <hcauchy (scale=0.1)>
## # ... with 25 more rows
```

Since $\eta$ is the precision of the beta distribution, it can be quite high. Let's set a less stringent prior:

```
m <- set_prior(m, normal(mean = 50, sd = 50), "^eta")
```

Since `medium` is a steady state compartment, let's set its `lambda` parameter to a dummy constant to save a bit of time during MCMC:

```
m <- set_prior(m, constant(0), "lambda_medium")
```

Let's run the model:

```
cache_file <- file.path("z-cache-case-study-li-2017-run.rds")
if (!file.exists(cache_file)) {
  run <- run_mcmc(m, iter = 2000)
  saveRDS(run, file = cache_file)
} else {
  run <- readRDS(cache_file)
}
plot(run)
```

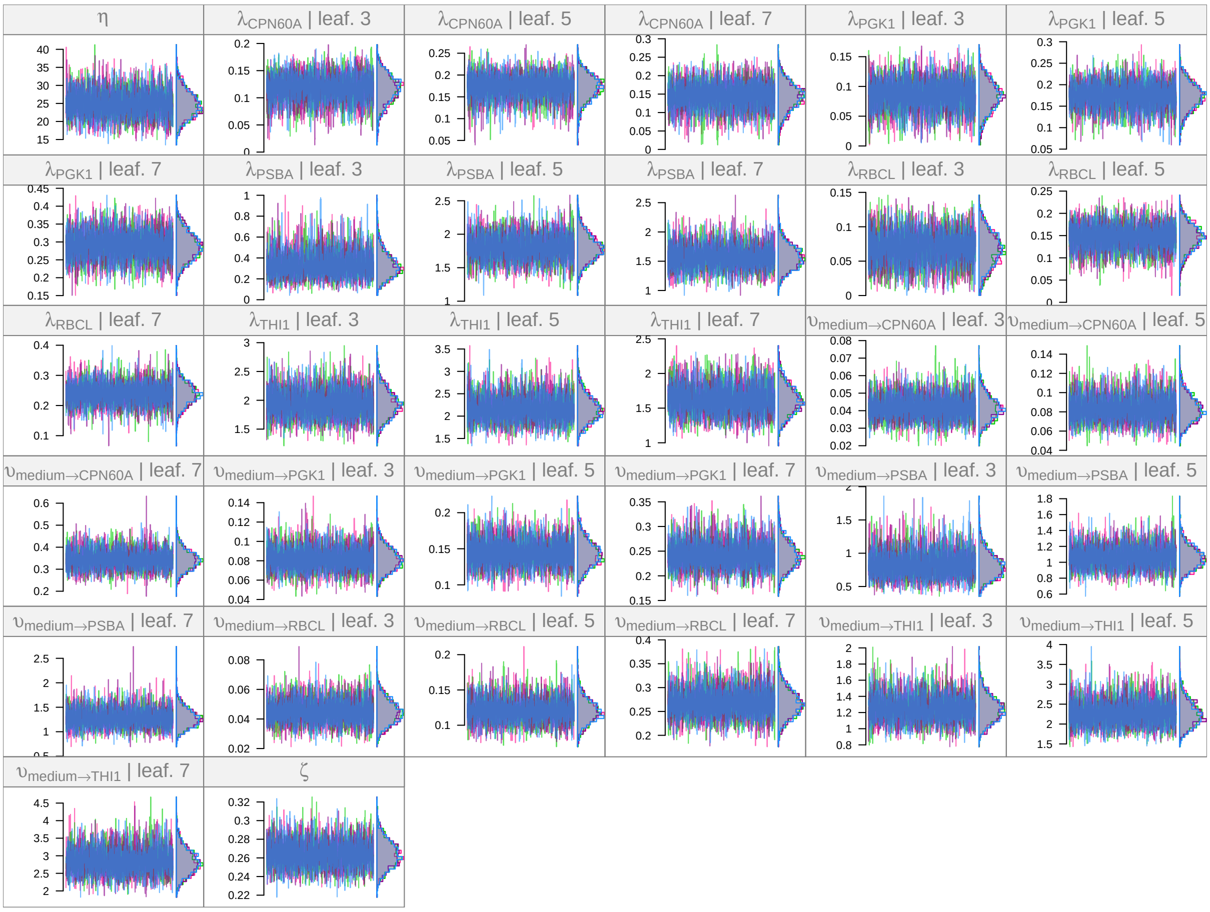


#### Posterior predictive checks

Let's see how the model fits the observations:

```r
cache_file <- file.path("z-cache-case-study-li-2017-pred.rds")
if (!file.exists(cache_file)) {
  pred <- m %>% predict(run, draws = 500, cores = n_cores)
  saveRDS(pred, file = cache_file)
} else {
  pred <- readRDS(cache_file)
}
```

```r
plot(pred, facet_row = "group", facet_col = "compartment", type = "prop",
     scale = "all", comps = c("CPN60A", "PGK1", "RBCL", "THI1", "PSBA"))
```

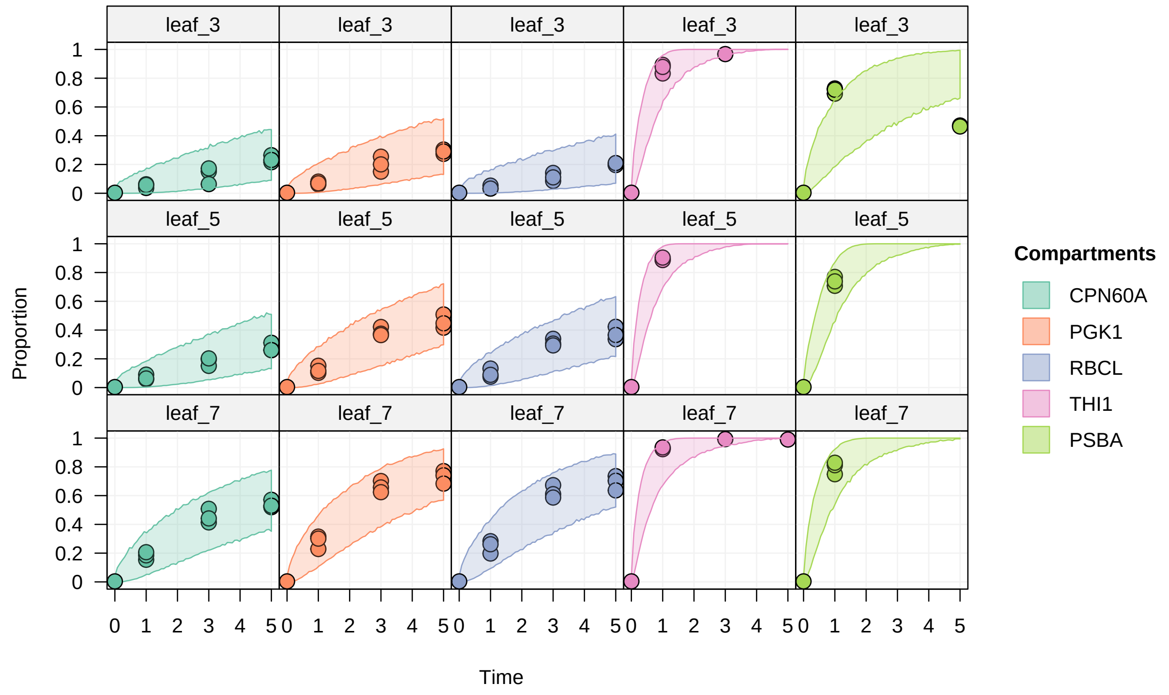


```r
plot(pred, facet_row = "group", facet_col = "compartment", type = "size",
     scale = "all", comps = c("CPN60A", "PGK1", "RBCL", "THI1", "PSBA"))
```

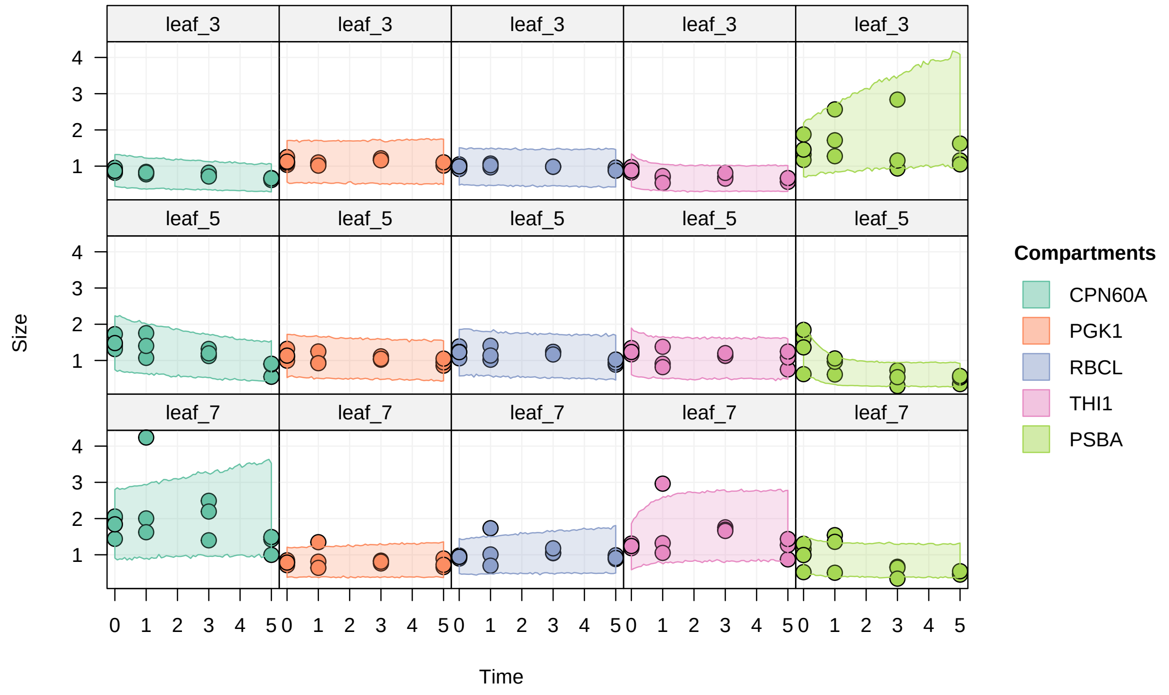


The model performs well to predict both labelled fractions and relative abundances.

#### Interpretation

One of the aim of the original paper was to estimate the turnover rate of as many proteins as possible in the three leaf types, and compare those rates across leaves and proteins.

Turnover rates are overall lower in leaf 3, and vert large differences exist between proteins. Large difference between leaves for a given protein:

```
library(ggplot2)
library(ggdist)

lambdas <- run %>% select("lambda") %>%
  tidy_mcmc() %>%
  mutate(mcmc.parameters = map(mcmc.parameters, enframe)) %>%
  pull(mcmc.parameters) %>%
  bind_rows() %>%
  separate(name, sep = "[|]", into = c("param", "leaf")) %>%
  mutate(prot = substr(param, 8, nchar(param)))

lambdas %>% group_by(prot, leaf) %>%
  median_qi(value, .width = c(0.8, 0.95)) %>%
  ggplot(aes(x = leaf, y = value, ymin = .lower, ymax = .upper)) +
  facet_grid(. ~ prot) +
  geom_pointinterval() +
  coord_trans(y = "log10") +
  labs(y = "lambda (day-1)")
```

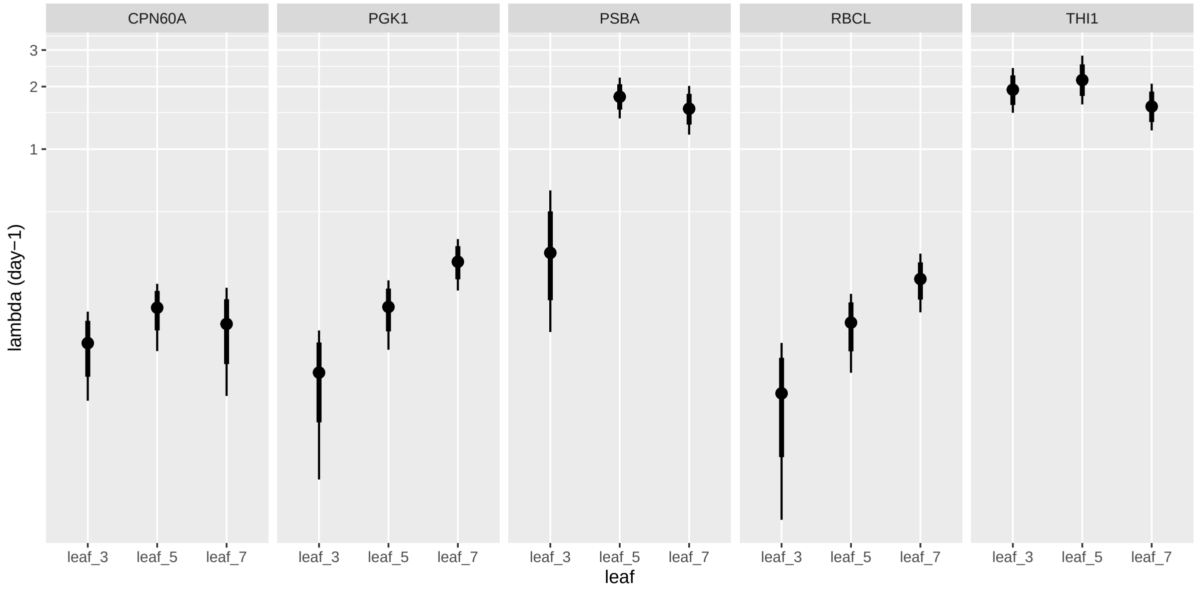


Conversely, the turnover times (calculated as the inverse of the turnover rates) range from less than 12 hours to tens of days, depending on the protein and on the leaf:

```
totimes <- lambdas %>% mutate(value = 1/value)

totimes %>% group_by(prot, leaf) %>%
  median_qi(value, .width = c(0.8, 0.95)) %>%
  ggplot(aes(x = leaf, y = value, ymin = .lower, ymax = .upper)) +
  facet_grid(. ~ prot) +
  geom_pointinterval() +
  coord_trans(y = "log10") +
  scale_y_continuous(breaks = c(1, 3, 5, 10, 20, 40, 60)) +
  labs(y = "turnover time (day)")
```

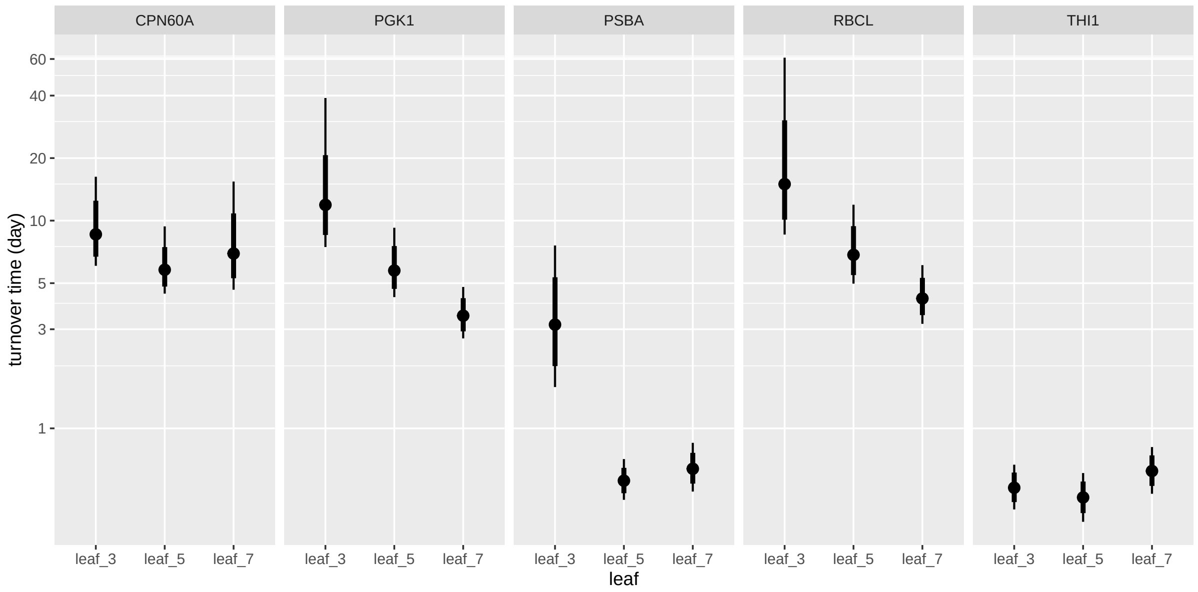


#### Sankey plots

We can calculate the flows from the model output. Note that those flows are not in absolute quantity of N since the protein pool size used here are the relative abundance compared to a reference sample. Those values could be converted to absolute values using the PaxDB which provides estimates of protein abundances for Arabidopsis (cf. the original paper).

Let's calculate the estimated average flows over the time of the experiment:

```
cache_file <- file.path("z-cache-case-study-li-2017-flows.rds")
if (!file.exists(cache_file)) {
  flows <- tidy_flows(m, run, n = 500)
  saveRDS(flows, file = cache_file)
} else {
  flows <- readRDS(cache_file)
}
```

Let's draw a Sankey plot for the 3rd and 7th leaves (note that the scaling is different between the two panels so they are not directly comparable, but the relative scalings of the flows and of the turnover times within each panel are correct):

```
library(grid)
grid.newpage()

vp <- viewport(layout = grid.layout(ncol = 2))
pushViewport(vp)

pushViewport(viewport(layout.pos.col = 1))
flows_3 <- flows %>% filter_by_group(leaf_id == "leaf_3")
quick_sankey(flows_3, node_s = "roundsquare", edge_f = 0.5, new = FALSE)
popViewport()

pushViewport(viewport(layout.pos.col = 2))
flows_7 <- flows %>% filter_by_group(leaf_id == "leaf_7")
quick_sankey(flows_7, node_s = "roundsquare", edge_f = 0.5, new = FALSE)
popViewport()
```

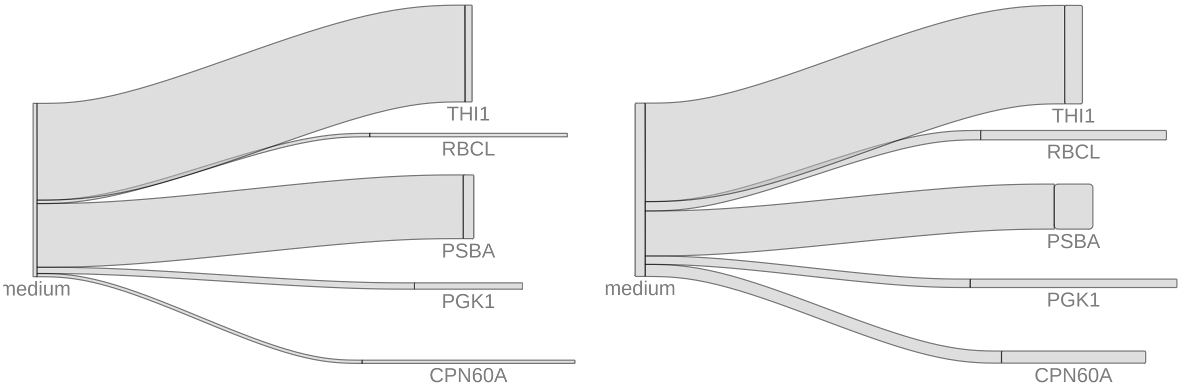
THI1
RBCL
PSBA
PGK1
CPN60A
medium
THI1
RBCL
PSBA
PGK1
CPN60A
medium

### Case study 2: Phosphorus uptake in eelgrass (McRoy and Barsdate 1970)

This case study models phosphorus uptake at the organism level in eelgrass from a dataset adapted from the following article:

- McRoy, C. Peter, and Robert J. Barsdate. "Phosphate Absorption in Eelgrass1." Limnology and Oceanography 15, no. 1 (January 1, 1970): 6–13. https://doi.org/10.4319/lo.1970.15.1.0006.

#### Biological question and experimental setup

Eelgrass can be present in dense populations in shallow water areas, and can have an important role in nutrients cycles in the ecosystems it is present in.

The McRoy 1970 study examines the uptake of phosphorus by eelgrass plants from the surrounding water or sediment and its transfer in the plant tissues. What is the preferred uptake pathway for phosphorus (leaves or roots)? How fast is it exchanged across plant tissues? Do the uptake processes depend on light conditions?

The experimental setup consists in individual eelgrass plant kept in closed jars, with a watertight septum separating the upper and the lower compartments. The upper compartment contains the leaves and stems of the plant, while the lower compartment contais the roots and rhizome. 32P is added as phosphate to either the upper or lower water compartment, and jars are kept in either light or dark conditions. At several time points over the course of two days, individual plants are taken from the experiment and 32P abundance in the water and in the plant tissues is quantified by measuring radioactive decay (cpm/mg of dry material).

#### Modelling

##### Data preparation

```
library(tidyverse)
library(isotracer)
```

The data is available in `eelgrass`:

```
eelgrass
```

```
## # A tibble: 83 x 7
##    light_treatment addition_site compartment time_min n_32P_per_mg mass_mg      n_32P
##    <chr>           <chr>         <chr>          <int>        <dbl>   <dbl>      <dbl>
##  1 dark            upper         leaves_stem        1     8163097.      46     3.76e8
##  2 dark            upper         leaves_stem      120    17135877.      46     7.88e8
##  3 dark            upper         leaves_stem      600    28156120.      46     1.30e9
##  4 dark            upper         leaves_stem     1500   157367757.      46     7.24e9
##  5 dark            upper         leaves_stem     3000    97868573.      46     4.50e9
##  6 dark            upper         lower_water        1        3474.   80000     2.78e8
##  7 dark            upper         lower_water      120        9332.   80000     7.47e8
##  8 dark            upper         lower_water      600       59509.   80000     4.76e9
##  9 dark            upper         lower_water     1500       18571.   80000     1.49e9
## 10 dark            upper         lower_water     3000        4448.   80000     3.56e8
## # ... with 73 more rows
```

In this experiment, we only have observations for the total amount of 32P per compartment (we do not measure the amount of 31P). The `isotracer` model will be provided with proportions of 1 for all measurements, and we will not estimate $\eta$ in the model:

```
eelgrass <- eelgrass %>% mutate(prop = 1)
```

We modify the data so that the time is given in days (this will make specifying the half-life of 32P more straightforward since it is usually given in days). We also modify the units for the abundance of 32P: instead of atom counts we will use $10^6$ atoms to make things more readable:

```
eelgrass <- eelgrass %>%
  mutate(time_day = time_min / (24 * 60),
         n_1e6_32P = n_32P / 1e6)
```

We keep only the columns we will need for the modelling:

```
eelgrass <- eelgrass %>%
  select(compartment, time_day, n_1e6_32P, prop, light_treatment, addition_site)
```

This is our table ready for the model specification:

```
eelgrass
```

```
## # A tibble: 83 x 6
##    compartment time_day n_1e6_32P  prop light_treatment addition_site
##    <chr>          <dbl>     <dbl> <dbl> <chr>           <chr>
##  1 leaves_stem 0.000694      376.     1 dark            upper
##  2 leaves_stem 0.0833        788.     1 dark            upper
##  3 leaves_stem 0.417        1295.     1 dark            upper
##  4 leaves_stem 1.04         7239.     1 dark            upper
##  5 leaves_stem 2.08         4502.     1 dark            upper
##  6 lower_water 0.000694      278.     1 dark            upper
##  7 lower_water 0.0833        747.     1 dark            upper
##  8 lower_water 0.417        4761.     1 dark            upper
##  9 lower_water 1.04         1486.     1 dark            upper
## 10 lower_water 2.08          356.     1 dark            upper
## # ... with 73 more rows
```

##### Model fitting

There are two treatments with two levels each (light/dark and 32P added in upper/lower water compartment) and the treatments are crossed.

We will assume that the rates defining the phosphorus flows do not depend on the compartment where 32P is added, but that the light conditions can have an effect on the rates. In other words: light condition is a covariate for the rates, while addition compartment is simply a replication unit.

Let's consider the following network topology:

```
m <- new_networkModel() %>%
  set_topo("upper_water -> leaves_stem -> roots_rhizome",
           "lower_water -> roots_rhizome -> leaves_stem")
ggtopo(m)
```

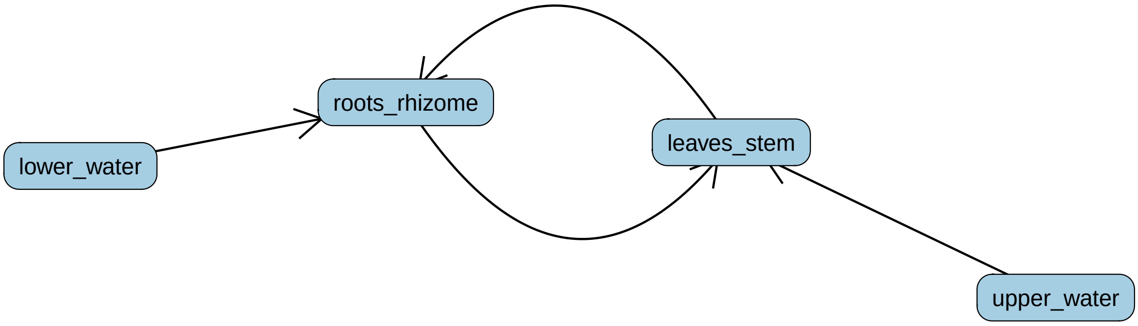


Note that to make the model more amenable, we have neglected the release of phosphorus by the plant into the water (but this is actually an important biological process!).

(When including release from the plant, the model estimates some very high exchange rates!)

32P is a radioactive compound and undergoes decay, which has to be taken into account in the model given the experiment time scale. The half-life of 32P is about 14 days:

```
m <- m %>% set_half_life(14.268)
```

Let's separate the initial conditions from the later observations:

```
init <- filter(eelgrass, time_day < 0.01)
obs <- filter(eelgrass, time_day > 0.01)
```

We add those initial conditions and the observations to the model. Note that we specify that the experimental data is grouped by treatment (light condition and location of tracer addition):

```
m <- m %>%
  set_init(init, comp = "compartment", size = "n_1e6_32P", prop = "prop",
           group_by = c("light_treatment", "addition_site")) %>%
  set_obs(obs, time = "time_day")
```

The default parameters of the model are:

```
params(m)
```

```
##  [1] "eta"                                  "lambda_leaves_stem"
##  [3] "lambda_lower_water"                   "lambda_roots_rhizome"
##  [5] "lambda_upper_water"                   "upsilon_leaves_stem_to_roots_rhizome"
##  [7] "upsilon_lower_water_to_roots_rhizome" "upsilon_roots_rhizome_to_leaves_stem"
##  [9] "upsilon_upper_water_to_leaves_stem"   "zeta"
```

We assume that no 32P exits the system (except what is lost through radioactive decay), so we set all `lambda` parameters to constant 0:

```
m <- m %>% set_prior(constant(0), "lambda")
```

Since $\eta$ has no role in this model (only "marked" tracer is observed), we just set it to a constant dummy value:

```
m <- m %>% set_prior(constant(1), "^eta")
```

To avoid the model exploring very large and unlikely values for the `upsilon` rates and for the `zeta` coefficient of variation of observed pool sizes, we give them a wide but bounded uniform prior over (0, 10):

```
m <- m %>%
  set_prior(uniform(0, 10), "upsilon") %>%
  set_prior(uniform(0, 10), "zeta")
```

Priors are set, but we still need to specify that we want to use the light condition as a covariate for the uptake rates:

```
m <- m %>% add_covariates(upsilon ~ light_treatment)
```

We are now ready to fit the model:

```
fit <- run_mcmc(m)
```

How do the chains look like?

```
plot(fit)
```

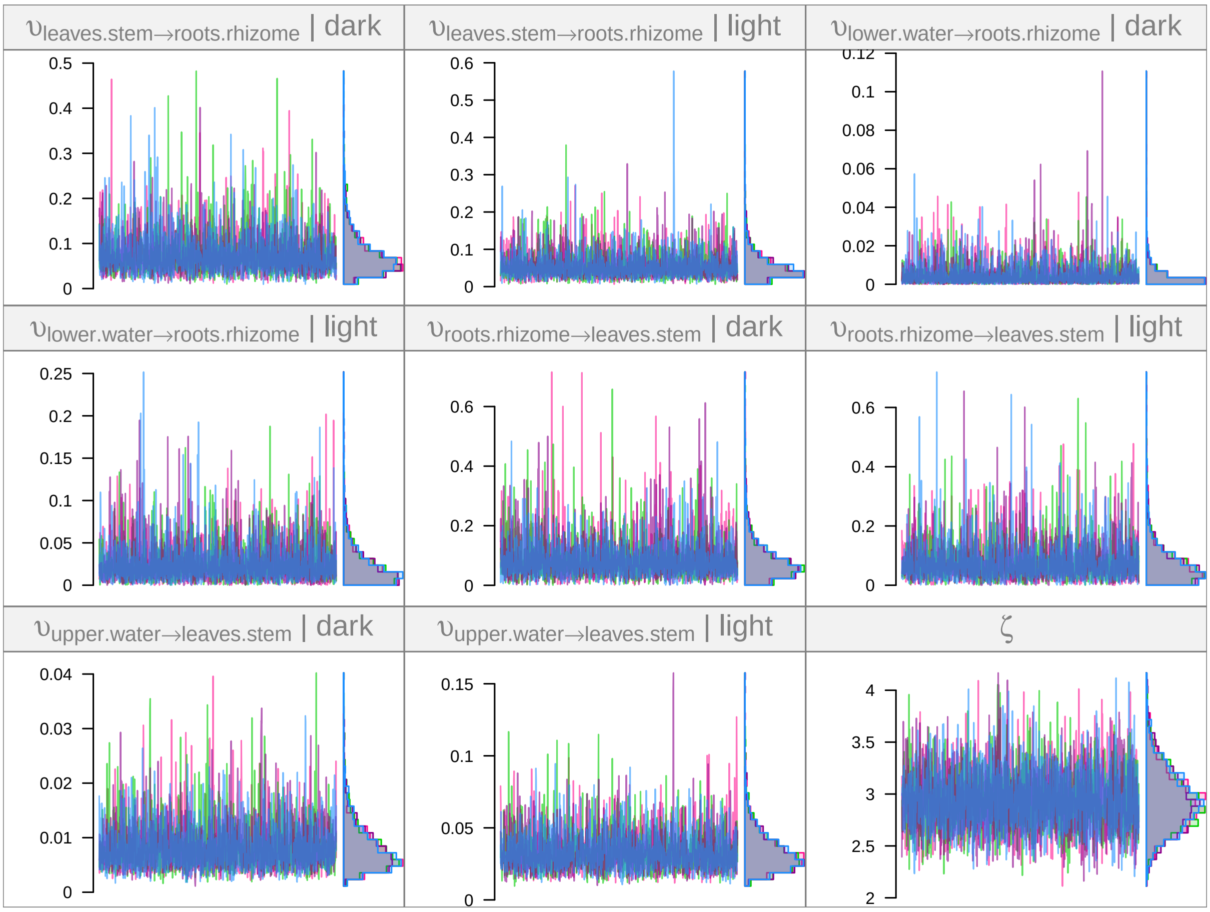


The chains look fine. How can we interpret our parameter estimates now?

#### Interpretation

The most striking result is that the estimated $\zeta$ value is extremely large: around 3! $\zeta$ is the estimated coefficient of variation for the pool size, so this means that the standard deviation of the observed sizes is about 3 times the mean expected size for any given time point! This is extremely large, but it is consistent with the data, which does contain ups and downs:

```
ggplot(eelgrass, aes(x = time_day, y = n_1e6_32P)) +
  geom_line(aes(col = compartment)) +
  facet_wrap(~ light_treatment + addition_site) +
  coord_trans(y = "log10")
```

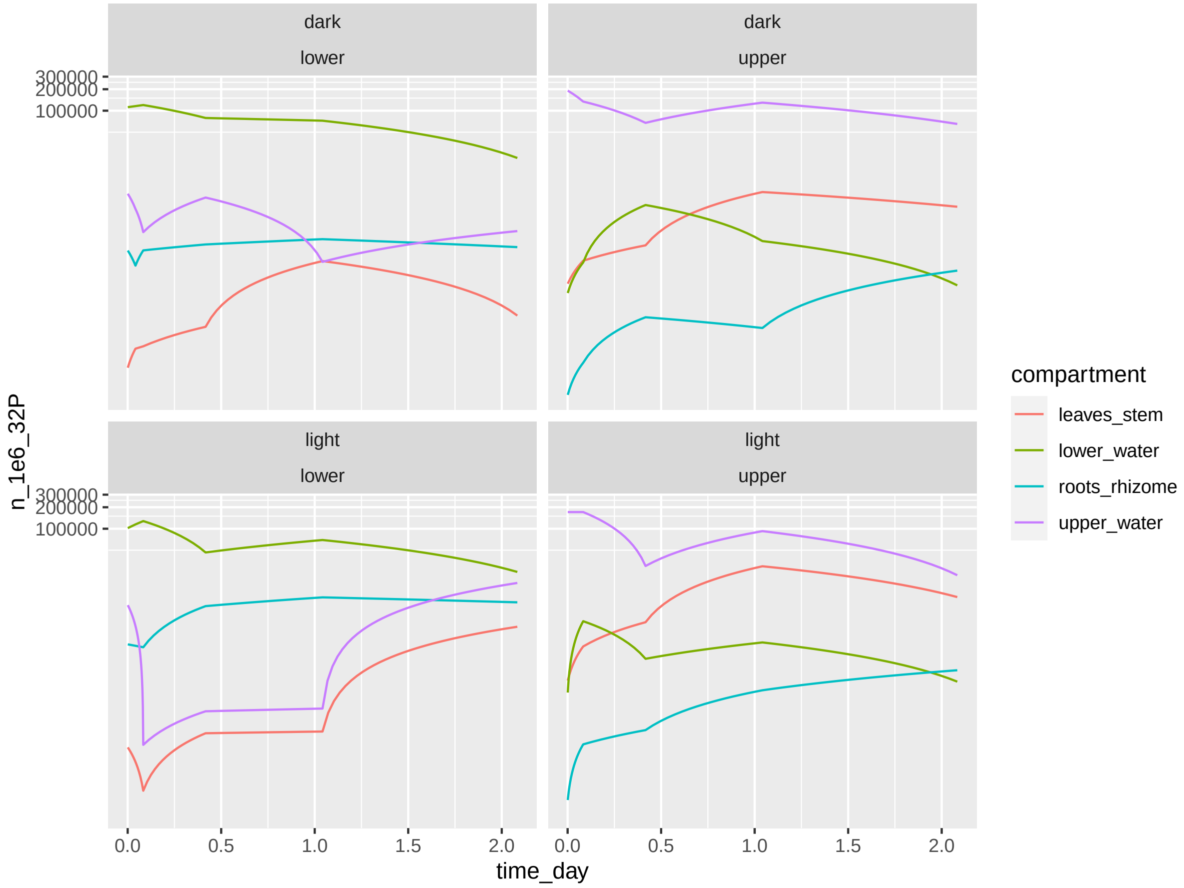


In the original article, authors explain this variability by the fact than distinct individual plants are sampled at each time point (to double-check). This is compounded by approximations we made when processing the data from the original paper:

- We pooled the 32P cpm/mg data from several tissues together and use their average value to reconciliate the pooling of the biomass data (given for e.g. “leaves and stem” together in the original paper) and the 32P data (given separately for leaf tip, middle and base and stem in the original paper)
- The reported data for each time point was in cpm/mg dry weight, and we use a single mean value of each tissue total dry weight per treatment to convert those values into 32P/compartment for the purpose of this case study.

Such a large $\zeta$ value is likely to impact the quality of the posterior predictions:

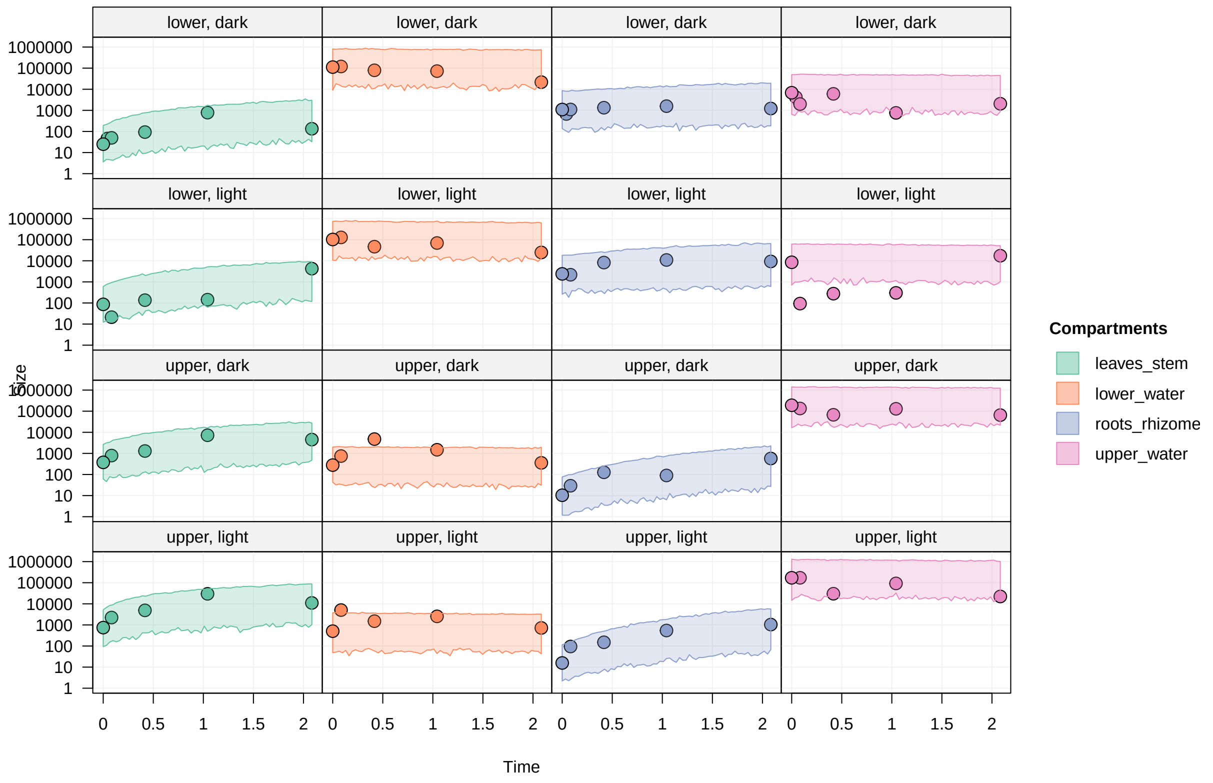


```
pred <- predict(m, fit, probs = 0.95, draws = 200)
plot(pred, facet_row = "group", facet_column = "compartment", log = TRUE, type = "size", scale = "al
```

As seen above, the predicted intervals for the trajectories of the pool sizes are extremely wide (note that the y scale is logarithmic). Hopefully, a dataset where total tissue dry weight is estimated at the same time as cpm/mg dry weight would allow to reduce a lot the uncertainty in those predicted trajectories.

Can we still say something about the parameter posteriors? Let's try. What are again the parameters we estimated?

```
varnames(fit)
```

```
## [1] "upsilon_leaves_stem_to_roots_rhizome|dark"
## [2] "upsilon_leaves_stem_to_roots_rhizome|light"
## [3] "upsilon_lower_water_to_roots_rhizome|dark"
## [4] "upsilon_lower_water_to_roots_rhizome|light"
## [5] "upsilon_roots_rhizome_to_leaves_stem|dark"
## [6] "upsilon_roots_rhizome_to_leaves_stem|light"
## [7] "upsilon_upper_water_to_leaves_stem|dark"
## [8] "upsilon_upper_water_to_leaves_stem|light"
## [9] "zeta"
```

One interesting question is whether the roots incorporate phosphate with the same efficiency as the leaves. Let's build a derived parameter calculating the ratio between the two `upsilon` rates in light condition:

```
z <- (fit[, "upsilon_lower_water_to_roots_rhizome|light"] /
      fit[, "upsilon_upper_water_to_leaves_stem|light"])
summary(z)$quantiles
```

```
##       2.5%        25%        50%        75%      97.5%
## 0.03354928 0.25964846 0.56776971 1.10555385 3.41674644
```

The uncertainty is too large to say anything in this case, even though it looks like the `upsilon` rate might be smaller for the roots than for the leaves. Can we ask another question?

What about the difference between light and dark conditions? For example, what is the effect of light on the 32P uptake by the leaves:

```
z <- (fit[, "upsilon_upper_water_to_leaves_stem|light"] /
      fit[, "upsilon_upper_water_to_leaves_stem|dark"])
summary(z)$quantiles
```

```
##      2.5%       25%       50%       75%     97.5%
##  1.339877  2.817730  4.271373  6.301579 13.165118
```

In this case, even with the large uncertainty in our estimates, we can rely say that the leaves incorporates 32P faster in light condition (mean estimate: about 5x faster).

Does light have an effect on the 32P uptake by the roots?

```
z <- (fit[, "upsilon_lower_water_to_roots_rhizome|light"] /
      fit[, "upsilon_lower_water_to_roots_rhizome|dark"])
summary(z)$quantiles
```

```
##        2.5%         25%         50%         75%       97.5%
##   0.2327051   2.2644152   6.6923988  20.4018232 263.8261930
```

Here again, the uncertainty is massive, but overall it looks like 32P uptake at the roots is probably faster under light conditions.

##### Sankey plot

We can build a Sankey plot representation of the 32P flows in this experiment, while keeping in mind that the uncertainties were very large and that the Sankey plot will only present average flow values.

To estimate flows for a situation where there is as much 32P in the water column and in the sediment, we will create some new initial conditions where there is as much 32P in the upper and lower water compartments (and the leaves and roots do not have any 32P) (to double-check: does this reflect a biological reality in the field?):

```
init2 <- tibble(compartment = c("lower_water", "upper_water", "roots_rhizome",
                                "leaves_stem"),
                n_1e6_32P = c(1e5, 1e5, 0, 0),
                prop = 1)
# Create a `m2` model with the adjusted initial conditions
m2 <- m %>%
  set_init(init2, comp = "compartment", size = "n_1e6_32P", prop = "prop")
```

Now we can estimate the flows given those initial conditions and the parameter posteriors:

```
flows <- tidy_flows(m2, fit, n = 200)
flows
```

```
## # A tibble: 800 x 5
##    group     mcmc.chain mcmc.iteration mcmc.parameters flows
##  * <list>         <int>          <int> <list>          <list>
##  1 <chr [2]>          3            678 <dbl [14]>      <grouped_df [8 x 3]>
##  2 <chr [2]>          4            109 <dbl [14]>      <grouped_df [8 x 3]>
##  3 <chr [2]>          2            162 <dbl [14]>      <grouped_df [8 x 3]>
##  4 <chr [2]>          4            441 <dbl [14]>      <grouped_df [8 x 3]>
##  5 <chr [2]>          4             57 <dbl [14]>      <grouped_df [8 x 3]>
##  6 <chr [2]>          3            821 <dbl [14]>      <grouped_df [8 x 3]>
##  7 <chr [2]>          2            571 <dbl [14]>      <grouped_df [8 x 3]>
##  8 <chr [2]>          1             78 <dbl [14]>      <grouped_df [8 x 3]>
##  9 <chr [2]>          1            526 <dbl [14]>      <grouped_df [8 x 3]>
## 10 <chr [2]>          4            740 <dbl [14]>      <grouped_df [8 x 3]>
## # ... with 790 more rows
```

This flow table contains estimates for all the treatment groups. As a reminder, the treatments were:

```r
groups(m2)
```

```
## # A tibble: 4 x 2
##   addition_site light_treatment
##   <chr>         <chr>
## 1 upper         dark
## 2 lower         dark
## 3 upper         light
## 4 lower         light
```

Let's draw a Sankey plot showing the flows estimated for the light treatment. With the `quick_sankey()` function, average flow estimates over the whole posterior are shown. We will use flows estimated from the additions in the upper and lower water compartments (so the estimates should be proportional to overall flows in a situation where phosphate is present in the sediment and in the water column):

```r
flows_light <- flows %>% filter_by_group(light_treatment == "light")
quick_sankey(flows_light, node_s = "roundsquare", edge_f = 0.25)
```

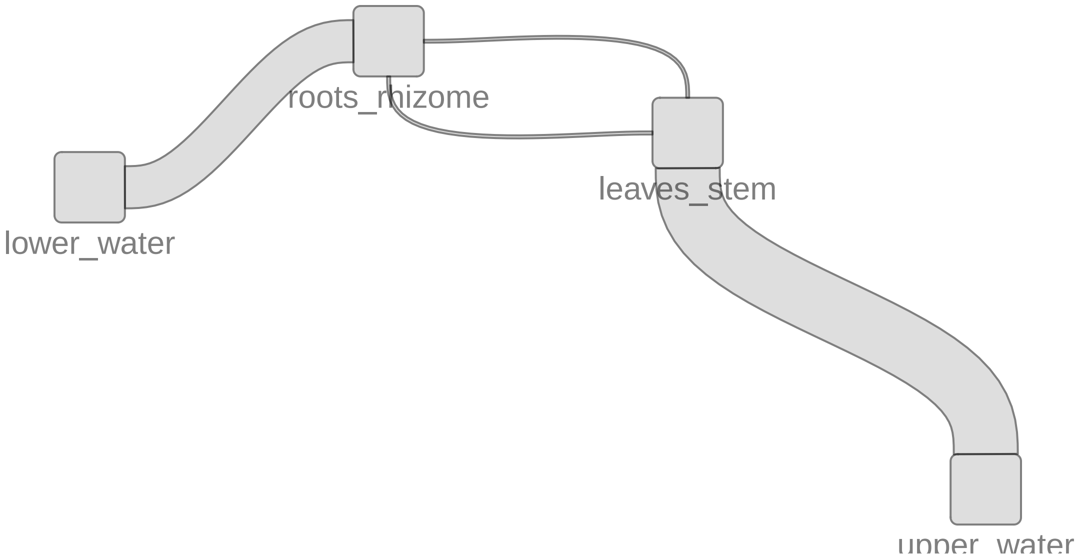

### Case study 3: Trinidadian streams (Collins et al. 2016)

This case study models nitrogen fluxes at the **ecosystem scale** from a dataset collected in Trinidadian mountain streams. The data is taken from the following article:

- Collins, S. M., S. A. Thomas, T. Heatherly, K. L. MacNeill, A. O.H.C. Leduc, A. López-Sepulcre, B. A. Lamphere, et al. **Fish Introductions and Light Modulate Food Web Fluxes in Tropical Streams: A Whole-Ecosystem Experimental Approach.** Ecology (2016) https://doi.org/10.1002/ecy.1530.

The goal is to quantify the nitrogen exchanges in the stream foodweb, from the dissolved nutrients ($NH_4^+$ and $NO_3^-$) to the top invertebrate predators. Using the **isotracer** package, we can:

- estimate those fluxes and the associated uncertainties
- test for the existence of suspected trophic links between some species of interest.

#### Experiment and data

In brief, $^{15}N$-enriched ammonium was dripped in two streams in Trinidad, and samples of the different foodweb compartments were taken during the drip and after the drip in several transects in each stream. The transects were located at different locations downstream of each drip. The drip phase lasted 10 days, and the post-drip phase lasted 30 days. Here is a glimpse of the dataset (we will use only the data from one stream for simplicity, but for a real analysis one could keep both streams and use stream identity as a covariate to estimate the stream effect on flows - see the case study about eelgrasses for an example of treatments comparison):

```
lalaja
```

```
## # A tibble: 223 x 6
##    stream transect   compartment time.days mgN.per.m2  prop15N
##    <chr>  <chr>      <chr>           <dbl>      <dbl>    <dbl>
##  1 UL     transect.1 FBOM                1      5657. NA
##  2 UL     transect.1 FBOM                1        NA   0.00372
##  3 UL     transect.1 FBOM                4        NA   0.00389
##  4 UL     transect.1 FBOM                7        NA   0.00434
##  5 UL     transect.1 FBOM               10        NA   0.00439
##  6 UL     transect.1 FBOM               14        NA   0.00428
##  7 UL     transect.1 FBOM               17        NA   0.00401
##  8 UL     transect.1 FBOM               20        NA   0.00424
##  9 UL     transect.1 FBOM               30        NA   0.00422
## 10 UL     transect.1 FBOM               40        NA   0.00390
## # ... with 213 more rows
```

- `stream` could be one of two locations (`LL` or `UL`), but only data from `UL` is used here.
- `transect` is nested within `stream`, and there are three transects per stream.
- `compartment` is a foodweb compartment.
- `time.days` is the sampling time.
- `mgN.per.m2` is the estimated size of the compartment.
- `prop15N` is the proportion of $^{15}N$ in the total nitrogen content of the compartment.

#### Building the model

##### Network topology

We will only model a simplified version of the foodweb described by Collins et al. to keep the model runtimes short. The foodweb model comprises:

- the two dissolved nutrients (ammonium and nitrate)

- two primary producers (epilithon, the layer of photosynthetic organisms growing on the stream bed, and FBOM, the fine benthic organic matter lying on the stream bed)
- three invertebrate grazers (*Petrophila*, *Psephenus* and *Tricorythodes*).
- one invertebrate predator (*Argia*), that preys on *Petrophila* and for which we would like to test another potential trophic link with *Psephenus*. We will test this extra trophic link later in the vignette.

We initialize the network model with the corresponding topology:

```
m <- new_networkModel() %>%
  set_topo("NH4, NO3 -> epi, FBOM", "epi -> petro, pseph", "FBOM -> tricor",
           "petro, tricor -> arg")
```

Since the inorganic nutrients are being constantly renewed by the stream flow, we will consider them in a steady state. We set both NH4 and NO3 to a steady state in the model:

```
m <- m %>% set_steady(c("NH4", "NO3"))
```

This is how our network look like at this stage:

```
ggtopo(m, "sugiyama")
```

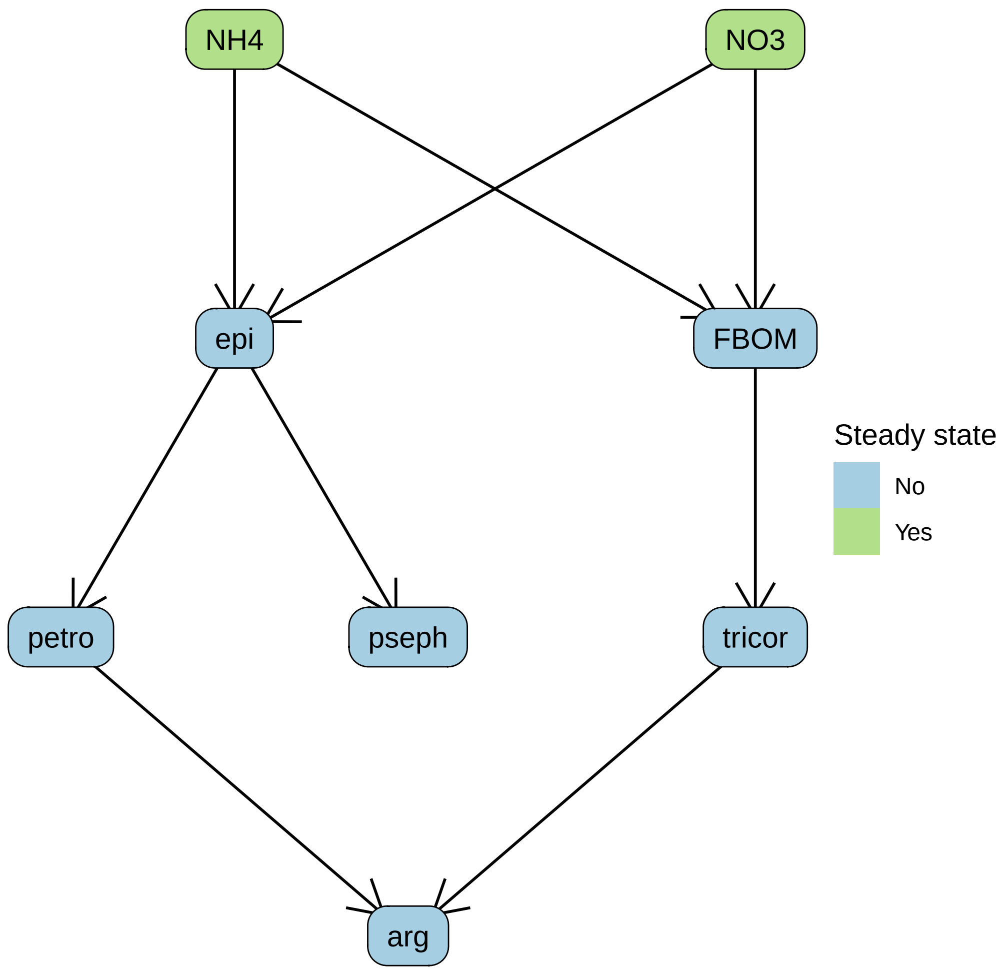

##### Initial conditions and observations

The drip of $^{15}N$-enriched ammonium was started at the very beginning of the experiment, and the initial conditions reflect the enrichment observed for NH4 and NO3 (only ammonium is dripped, but it is converted to nitrate quite rapidly and the experimental setup is thus equivalent to a drip phase for both ions).

The initial conditions are contained in a `tibble`:

```
inits
```

```
## # A tibble: 24 x 5
## # Groups:   stream, transect [3]
##    stream transect    compartment mgN.per.m2 prop15N
##    <chr>  <chr>       <chr>            <dbl>   <dbl>
##  1 UL     transect.1 NH4               0.300 0.0240
##  2 UL     transect.1 NO3              31.5   0.00397
##  3 UL     transect.2 NH4               0.300 0.0140
##  4 UL     transect.2 NO3              31.5   0.00407
##  5 UL     transect.3 NH4               0.300 0.00694
##  6 UL     transect.3 NO3              31.5   0.00413
##  7 UL     transect.1 arg               2.31  0.00366
##  8 UL     transect.2 arg               2.31  0.00366
##  9 UL     transect.3 arg               2.31  0.00366
## 10 UL     transect.1 epi             329.    0.00366
## # ... with 14 more rows
```

We add them to the model, specifying the grouping variable (`transect`, each transect being a different location downstream the drip station in the `UL` stream):

```
m <- m %>% set_init(inits, comp = "compartment", size = "mgN.per.m2", prop = "prop15N",
                    group_by = c("transect"))
```

The replication structure of the model can be examined using the `groups()` function:

```
groups(m)
```

```
## # A tibble: 3 x 1
##   transect
##   <chr>
## 1 transect.1
## 2 transect.2
## 3 transect.3
```

The next step is to add the observations made during the experiment:

```
m <- m %>% set_obs(trini, time = "time.days")
```

The enrichment data can be visualized with the `plot()` method:

```
# Note the log scale for better visualization
plot(m, facet_row = "group", facet_col = "compartment", type = "prop",
     scale = "all", log = TRUE,
     comps = c("NH4", "NO3", "epi", "FBOM", "petro", "pseph", "tricor", "arg"))
```

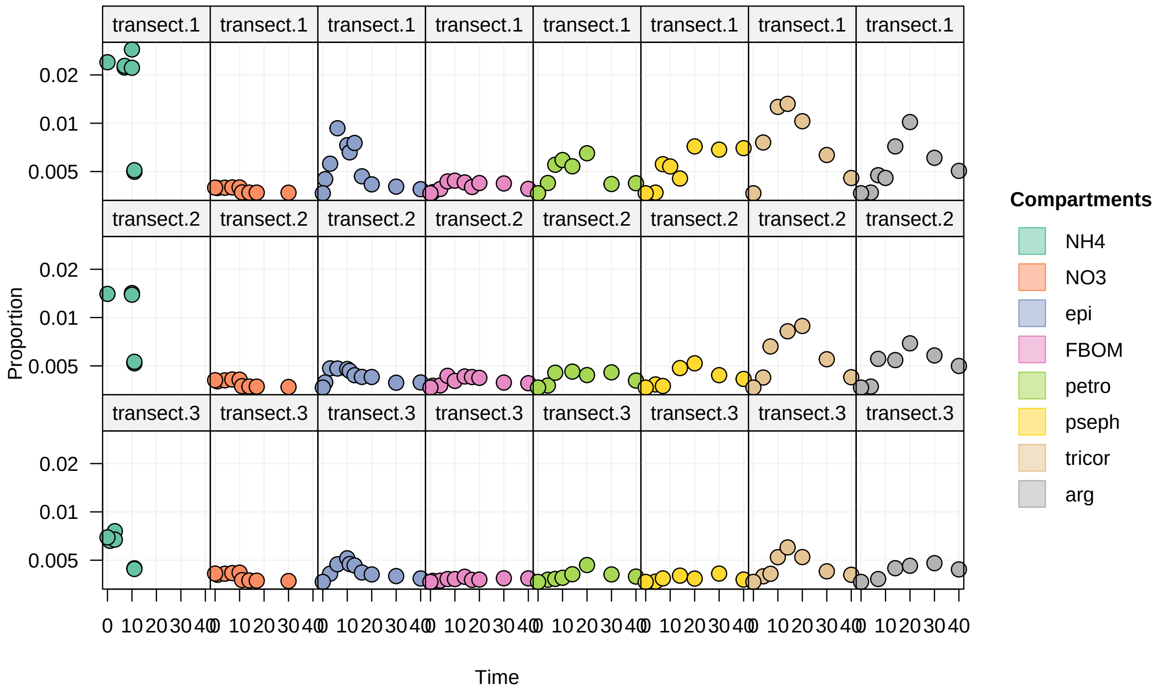


Interestingly, we see that one of the consumer, `tricor`, is over-enriched compared to its resources (`epi` and `FBOM`). This is due to the "bulk" nature of the epilithon and FBOM compartments.

##### Allowing for inhomogeneous compartments

Some compartment are sampled "in bulk" by the field scientist, but are really comprised of several sub-compartments, which can independently incorporate the marked tracer or be selectively fed upon by a consumer at the time scale of the experiment. For example, "epilithon" is really everything that we can be scratch and collect from the surface of a stream bed rock in the field, but it might actually be composed of an active fraction incorporating the dissolved nutrients (algae growing on the rock) and being selectively grazed by the invertebrate consumers, while the rest is a refractory fraction of sediment and debris which are not involved in the nutrient cycle at the scale of a few days.

To account for that in our model, we can allow for epilithon and FBOM to be split compartments with an active and a refractory portions:

```
m <- m %>% set_split(c("epi", "FBOM"))
```

For `epi` (and similarly for `FBOM`), this introduces a new parameter, `portion.act_epi`, which is used to split the initial biomass for the `epi` compartment into an active portion (involved in the network) and a refractory portion (isolated from the network, but used to predict observed size and proportion marked in the "bulk" epilithon).

##### Drip regime

The NH4 and NO3 quantities during the "on" phase of the drip (when the experiment starts) are already defined by (i) the fact that we set ammonium and nitrate to be in a steady state and (ii) the initial conditions. We will specify the quantities for the "off" phase by using the `add_pulse_event()` function to decrease the amount of $^{15}N$ in the ammomiun and nitrate compartments after day 10.

The `add_pulse_event()` function accepts either a single pulse specification, or bulk pulses defined in a table. We use the table syntax here:

```
pulses <- tribble(
  ~ stream,    ~ transect, ~ comp, ~ time, ~ qty_14N, ~ qty_15N,
      "UL",  "transect.1",  "NH4",     11,         0,  -0.00569,
```

```
      "UL",  "transect.2",  "NH4",     11,         0,  -0.00264,
      "UL",  "transect.3",  "NH4",     11,         0, -0.000726,
      "UL",  "transect.1",  "NO3",     11,         0,  -0.00851,
      "UL",  "transect.2",  "NO3",     11,         0,  -0.01118,
      "UL",  "transect.3",  "NO3",     11,         0,  -0.01244,
  )
```

```
m <- m %>% add_pulse_event(pulses = pulses, comp = "comp", time = "time",
                           unmarked = "qty_14N", marked = "qty_15N")
```

###### Setting the distribution family for observed sizes

Let's think a moment about the modelling of the observed compartment sizes. By default, the `isotracer` model assumes that observed compartment pool sizes are distributed following a normal distribution around the values expected from the calculated trajectories, with a coefficient of variation $\zeta$ shared by all compartments. This is fine when the compartments exhibit similar variation in their biomasses, but in the case of this ecosystem study they do not.

To illustrate this point, let's calculate the standard deviations and the corresponding coefficients of variation for the observed compartment sizes:

```
trini %>% select(compartment, mgN.per.m2) %>%
  na.omit() %>%
  group_by(compartment) %>%
  summarize(mean = mean(mgN.per.m2),
            sd = sd(mgN.per.m2),
            cv = sd / mean) %>%
  arrange(cv)
```

```
## # A tibble: 8 x 4
##   compartment    mean      sd    cv
##   <chr>         <dbl>   <dbl> <dbl>
## 1 NH4           0.300    0     0
## 2 NO3          31.5      0     0
## 3 epi         329.     115.   0.350
## 4 FBOM       5267.    2436.   0.463
## 5 pseph        14.4     10.7  0.745
## 6 arg           2.31     2.88 1.25
## 7 tricor        8.38    13.3  1.59
## 8 petro         1.40     2.48 1.77
```

The coefficients of variation are particularly large for the invertebrates, which is due to the patchiness of their distribution on the stream bed.

One way to handle this in our model is to allow for $\zeta$ to be compartment-specific. To make things easier for the model fit here, we will actually provide those compartment-specific $\zeta$ values as fixed parameters, but we could also let the model estimate them.

First, we specify that $\zeta$ depends on the compartment, and that it represents the standard deviation of the distributions:

```
m <- m %>% set_size_family("normal_sd", by_compartment = TRUE)
```

Then, we provide constant priors for those parameters based on the values calculated above:

```
m <- set_prior(m, constant(0.1), "zeta_NH4") %>%  # Dummy values for the steady
  set_prior(constant(0.1), "zeta_NO3") %>%        # state dissolved nutrients
  set_prior(constant(115), "zeta_epi") %>%
  set_prior(constant(2436), "zeta_FBOM") %>%
  set_prior(constant(10.7), "zeta_pseph") %>%
  set_prior(constant(2.88), "zeta_arg") %>%
```

```
  set_prior(constant(13.3), "zeta_tricor") %>%
  set_prior(constant(2.48), "zeta_petro")
```

#### Running the model

##### Setting priors

Before we run the model, let's have a look at the default priors used for the other parameters:

```
priors(m) %>% filter(!startsWith(in_model, "zeta"))
```

```
## # A tibble: 20 x 2
##    in_model                prior
##    <chr>                   <list>
##  1 eta                     <hcauchy (scale=0.1)>
##  2 lambda_arg              <hcauchy (scale=0.1)>
##  3 lambda_epi              <hcauchy (scale=0.1)>
##  4 lambda_FBOM             <hcauchy (scale=0.1)>
##  5 lambda_NH4              <hcauchy (scale=0.1)>
##  6 lambda_NO3              <hcauchy (scale=0.1)>
##  7 lambda_petro            <hcauchy (scale=0.1)>
##  8 lambda_pseph            <hcauchy (scale=0.1)>
##  9 lambda_tricor           <hcauchy (scale=0.1)>
## 10 portion.act_epi         <uniform (min=0,max=1)>
## 11 portion.act_FBOM        <uniform (min=0,max=1)>
## 12 upsilon_epi_to_petro    <hcauchy (scale=0.1)>
## 13 upsilon_epi_to_pseph    <hcauchy (scale=0.1)>
## 14 upsilon_FBOM_to_tricor  <hcauchy (scale=0.1)>
## 15 upsilon_NH4_to_epi      <hcauchy (scale=0.1)>
## 16 upsilon_NH4_to_FBOM     <hcauchy (scale=0.1)>
## 17 upsilon_NO3_to_epi      <hcauchy (scale=0.1)>
## 18 upsilon_NO3_to_FBOM     <hcauchy (scale=0.1)>
## 19 upsilon_petro_to_arg    <hcauchy (scale=0.1)>
## 20 upsilon_tricor_to_arg   <hcauchy (scale=0.1)>
```

Those default priors have to be adjusted to the data being modeled. Here, the time units are days, and so a half-Cauchy prior with a median of 0.1 means that the prior for the uptake rates and loss rates has a median equivalent to 10% of the N content of a given compartment being uptaken or lost in a day. This is reasonable for biological compartments, but it is too restrictive for dissolved nutrients: those are being constantly renewed by the stream flow, so we can expect much higher daily uptake rates from those.

Below we modify the priors for the uptakes from dissolved nutrients to allow for such higer daily uptake rates. In addtion, we also set their loss rate to a constant dummy value: since those compartments are in a steady state, we don't need the sampler to sample their loss rate.

```
m <- m %>%
  set_prior(hcauchy(5), "upsilon_N") %>%
  set_prior(constant(0), "lambda_N")
```

##### Running the model

We are ready to run the model fit:

```
run <- run_mcmc(m, iter = 2000)
plot(run)
```

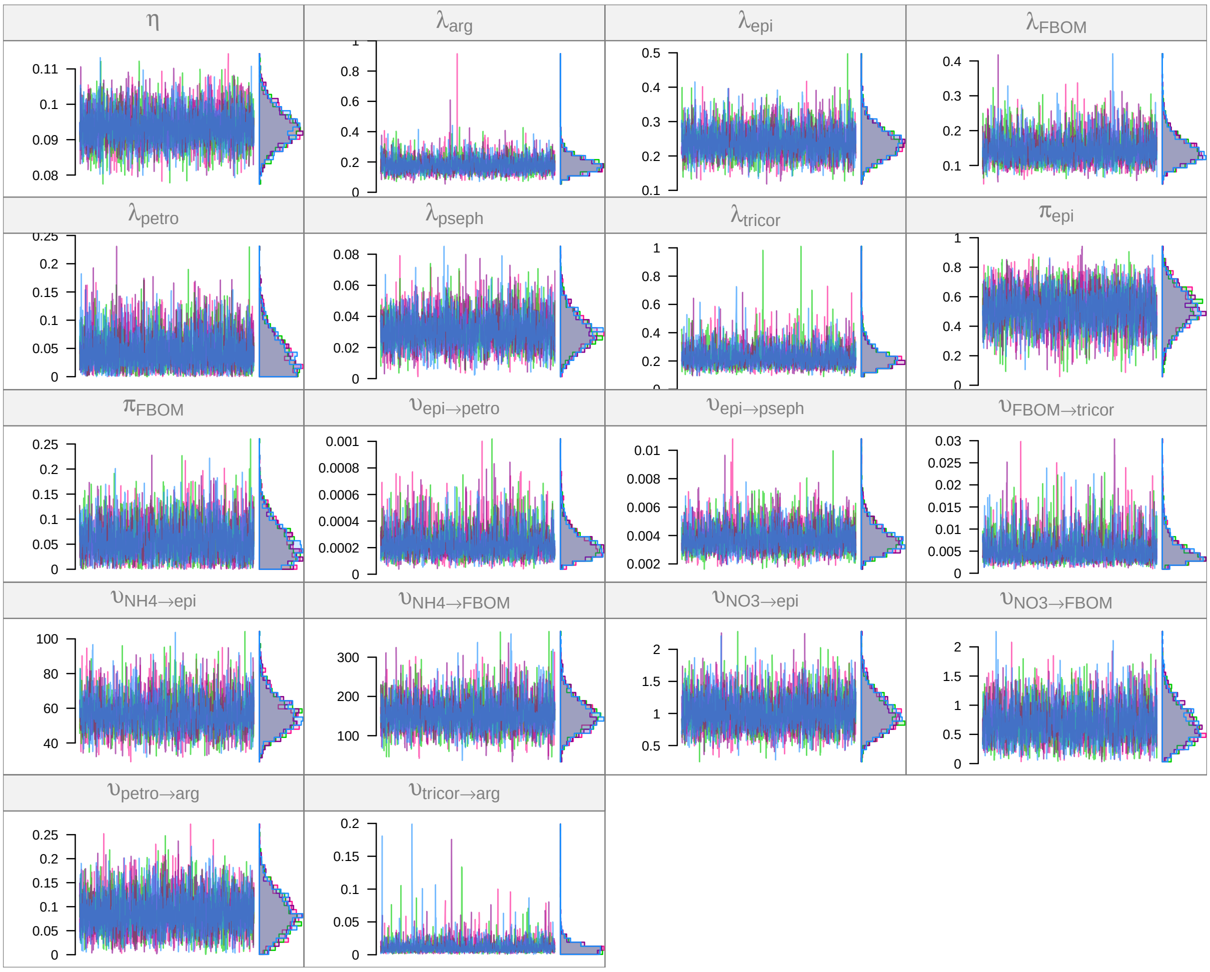


##### Posterior predictive fits

Once the fit is completed (and provided that no issues arised during the Stan run), we can calculate the posterior predicted trajectories and check if they match with the observed data:

```
pred <- predict(m, run, draws = 200)
```

```
plot(pred, facet_row = "group", facet_col = "compartment", type = "prop",
     scale = "all", log = TRUE,
     comps = c("epi", "FBOM", "petro", "pseph", "tricor", "arg"))
```

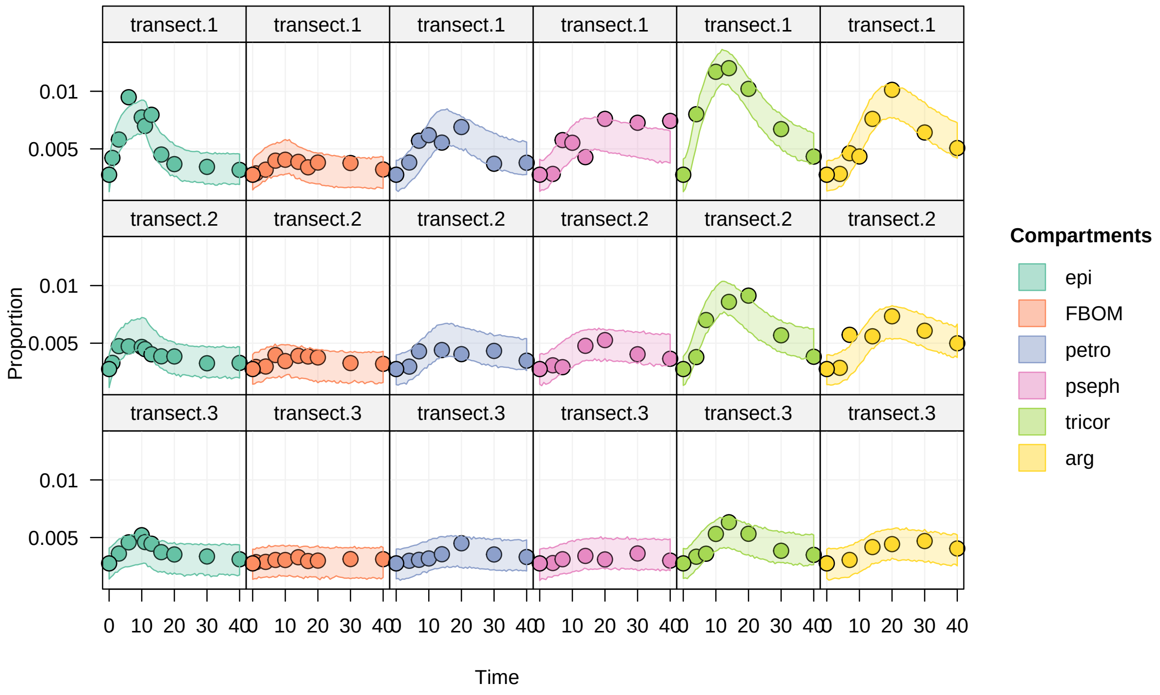


The quality of the fit looks quite good. We could use this fit to estimate flows between compartments and make quantitative conclusions from those results. We can also use this model as a base model to test for putative trophic link.

For example, one putative link is that *Argia* might feed on *Psephenus* in addition to feeding on *Petrophila* and *Tricorythodes*. We can build an run an alternative model with this link, and compare the two models.

#### Alternative model

The data used for this alternative model is the same, only the topology of the model is changed:

```
m2 <- m %>% set_topo("NH4, NO3 -> epi, FBOM", "epi -> petro, pseph",
                     "FBOM -> tricor", "petro, pseph, tricor -> arg")
```

Changing the topology resets steady and split compartments and priors. Let's apply the same settings as for the first model:

```
m2 <- m2 %>%
  set_steady(c("NH4", "NO3")) %>%
  set_split(c("epi", "FBOM")) %>%
  set_prior(constant(0.1), "zeta_NH4") %>%
  set_prior(constant(0.1), "zeta_NO3") %>%
  set_prior(constant(115), "zeta_epi") %>%
  set_prior(constant(2436), "zeta_FBOM") %>%
  set_prior(constant(10.7), "zeta_pseph") %>%
  set_prior(constant(2.88), "zeta_arg") %>%
  set_prior(constant(13.3), "zeta_tricor") %>%
  set_prior(constant(2.48), "zeta_petro") %>%
  set_prior(hcauchy(5), "upsilon_N") %>%
  set_prior(constant(0), "lambda_N")
```

The structure of this alternative model is:

```
ggtopo(m2, "sugiyama")
```

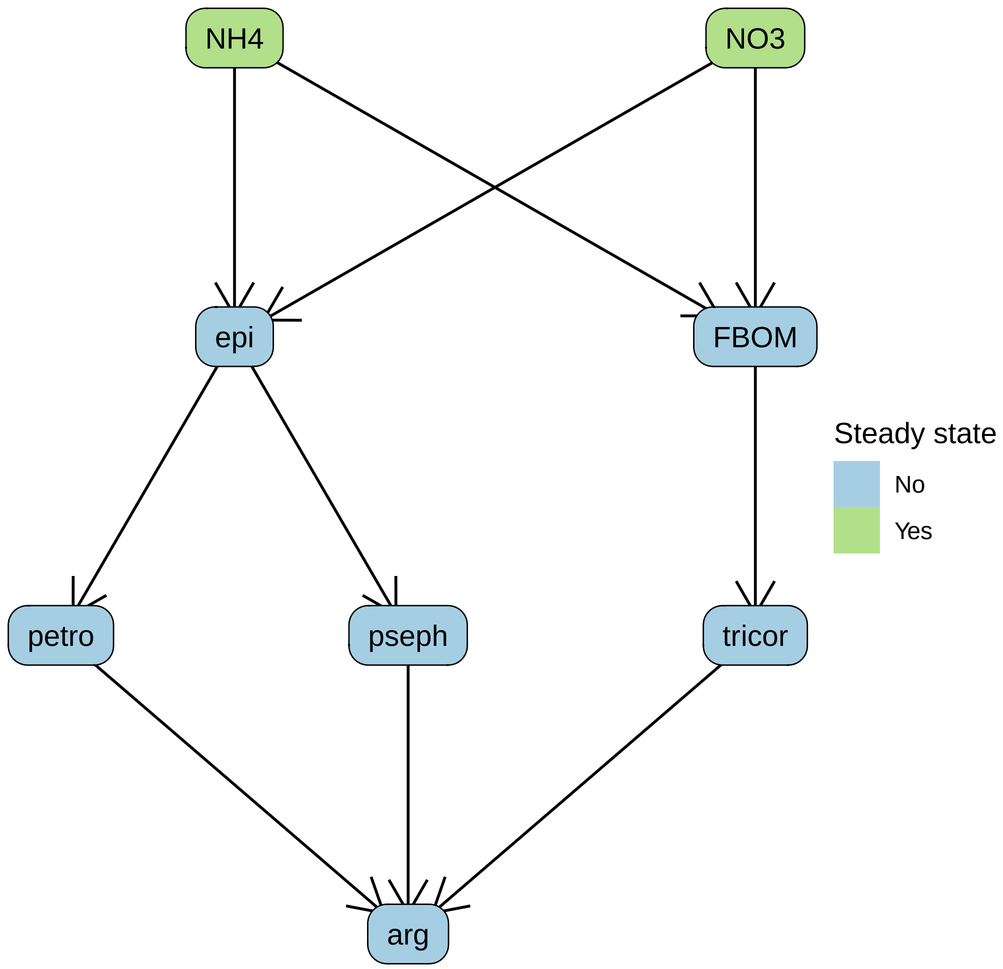


We run the model:

```
run2 <- run_mcmc(m2, iter = 2000)
plot(run2)
```

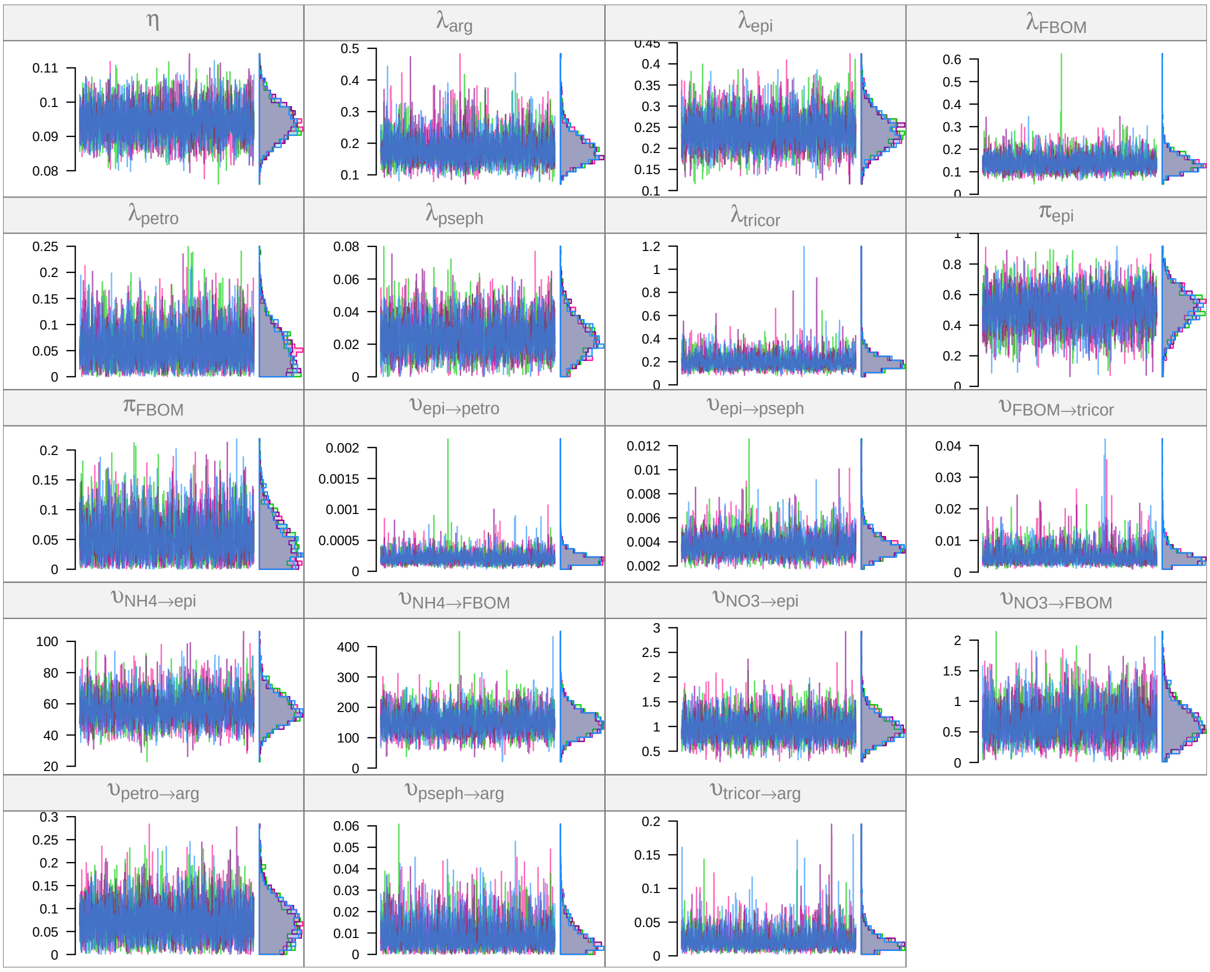

We check the model fit with a posterior predictive fit:

```
pred2 <- predict(m2, run2, draws = 200)
```

```
plot(pred2, facet_row = "group", facet_col = "compartment", type = "prop",
     scale = "all", log = TRUE,
     comps = c("epi", "FBOM", "petro", "pseph", "tricor", "arg"))
```

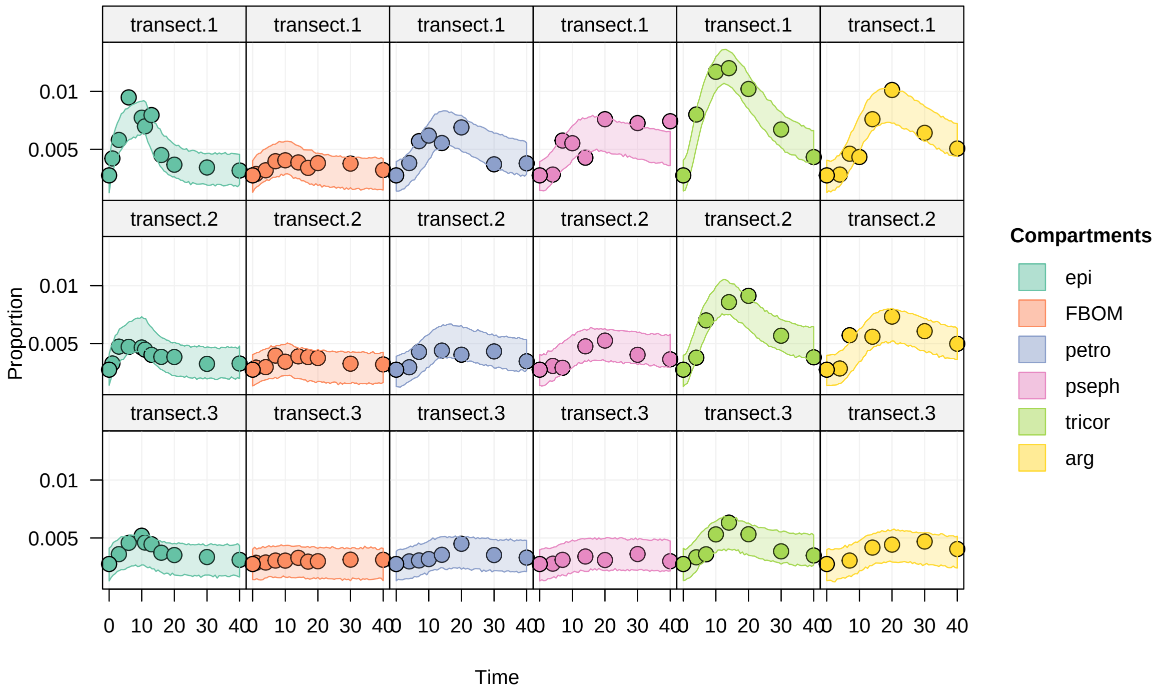


The fit looks equally good as before. Which model should we prefer?

#### Model comparison

We compare the two models by calculating their DIC and choosing the model with the lowest DIC.

Note that the use of DIC is not ideal here: DIC is valid when the posteriors have a multinormal distribution, which is not the case here givn that the parameters are bounded by zero and several exhibit skewness.

An alternative to DIC is to use LOO (leave-one-out validation). However, LOO requires independent data, and the data used here is a time series where data points are not independent. To check: WAIC?

Add refs!

We use the `dic()` function to calculate the model DIC values and the corresponding DIC weight:

```
dic(run, run2)
```

```
## # A tibble: 2 x 6
##   fit      Dbar    pD    DIC delta_DIC weight
##   <chr>   <dbl> <dbl>  <dbl>     <dbl>  <dbl>
## 1 run    -1821.  20.6 -1801.      0     0.796
## 2 run2   -1820.  21.6 -1798.      2.73  0.204
```

`run` is from the model fit without the trophic link `pseph -> arg`. According to the DIC, this model is slightly more likely than the alternative model with the extra trophic link. The evidence is not very strong though, as evidenced by the small $\Delta$DIC value and the DIC weights.

#### Sankey plot

Let's estimate the flows between compartments. Since this network has steady state sources, we can estimate flows at equilibrium:

```
flows <- tidy_flows(m, run, n = 500, steady_state = TRUE)
```

Sankey plot:

```r
f <- flows %>%
  select(flows) %>%
  unnest(flows) %>%
  na.omit() %>% # We drop the rows corresponding to `lambda` losses
  group_by(from, to) %>%
  summarize(flow = mean(average_flow),
            sd = sd(average_flow),
            cv = sd / flow)
```

```
## `summarise()` has grouped output by 'from'. You can override using the `.groups` argument.
```

```r
nodes <- inits %>%
  ungroup() %>%
  filter(transect == "transect.1") %>%
  select(comp = compartment, size = mgN.per.m2) %>%
  mutate(label = comp)
sankey(topo(m), nodes, f %>% select(from, to, flow) %>% rename(width = flow),
       edge_f = 0.2, layout = "left2right")
```

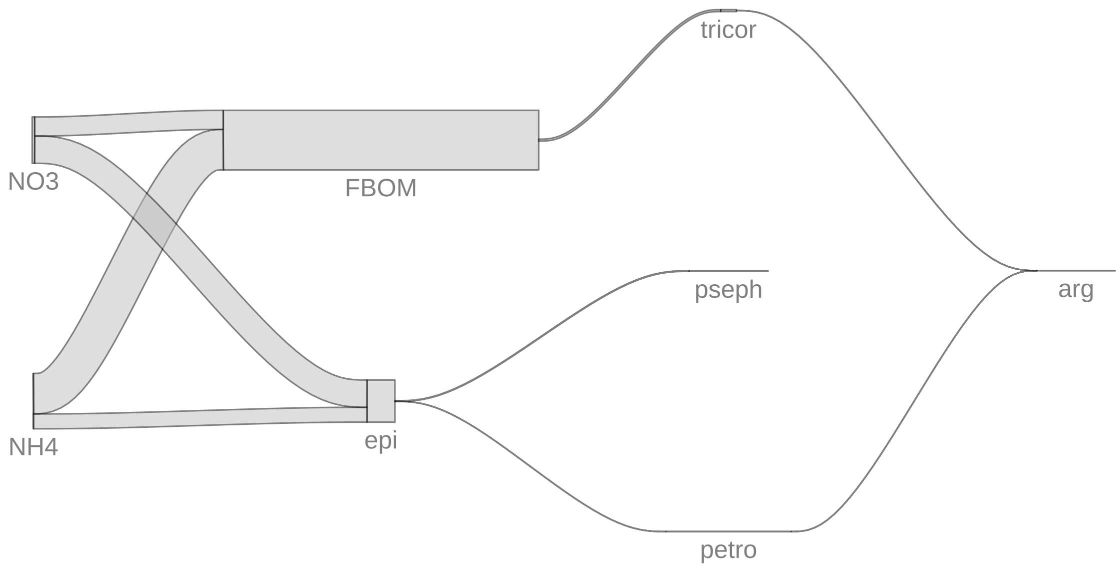


#### To go further

A heatmap of the correlation between parameter estimates can be a good way to visualize the strength of the interdependence between parameters:

```r
mcmc_heatmap(run)
```

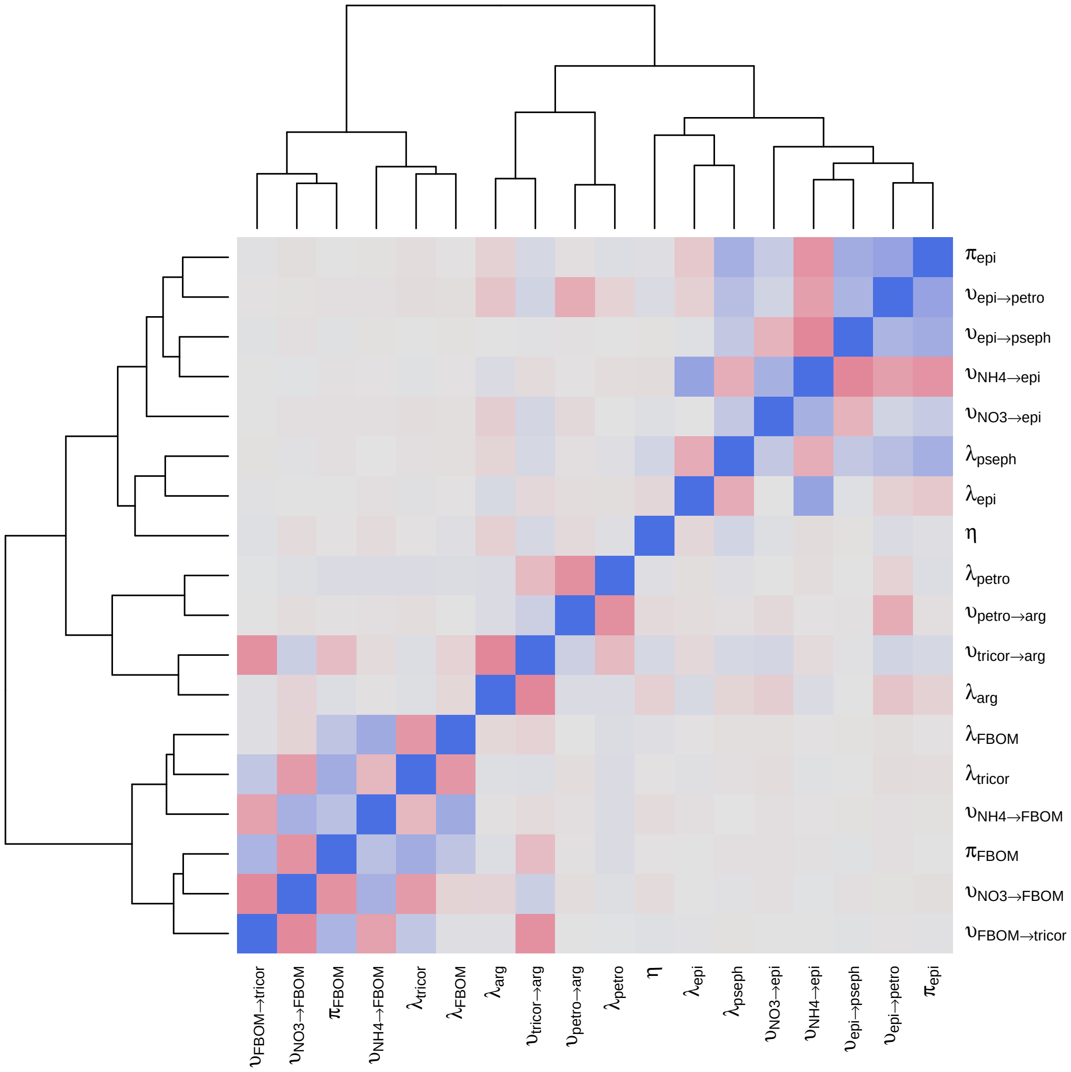

πepi
υepi→petro
υepi→pseph
υNH4→epi
υNO3→epi
λpseph
λepi
η
λpetro
υpetro→arg
υtricor→arg
λarg
λFBOM
λtricor
υNH4→FBOM
πFBOM
υNO3→FBOM
υFBOM→tricor

### Computing environment

```r
sessionInfo()
```

```
## R version 4.1.0 (2021-05-18)
## Platform: x86_64-pc-linux-gnu (64-bit)
## Running under: Ubuntu 18.04.5 LTS
##
## Matrix products: default
## BLAS:   /usr/lib/x86_64-linux-gnu/blas/libblas.so.3.7.1
## LAPACK: /usr/lib/x86_64-linux-gnu/lapack/liblapack.so.3.7.1
##
## locale:
##  [1] LC_CTYPE=en_US.UTF-8       LC_NUMERIC=C               LC_TIME=en_GB.UTF-8
##  [4] LC_COLLATE=en_US.UTF-8     LC_MONETARY=en_GB.UTF-8    LC_MESSAGES=en_US.UTF-8
##  [7] LC_PAPER=en_GB.UTF-8       LC_NAME=C                  LC_ADDRESS=C
## [10] LC_TELEPHONE=C             LC_MEASUREMENT=en_GB.UTF-8 LC_IDENTIFICATION=C
##
## attached base packages:
## [1] grid      stats     graphics  grDevices utils     datasets  methods   base
##
## other attached packages:
##  [1] ggdist_3.0.0        magrittr_2.0.1      forcats_0.5.1
##  [4] stringr_1.4.0       dplyr_1.0.7         purrr_0.3.4
##  [7] readr_2.0.0         tidyr_1.1.3         tibble_3.1.3
## [10] ggplot2_3.3.5       tidyverse_1.3.1     isotracer_0.7.6.9000
##
## loaded via a namespace (and not attached):
##  [1] matrixStats_0.60.0  fs_1.5.0            bit64_4.0.5
##  [4] lubridate_1.7.10    httr_1.4.2          rstan_2.21.2
##  [7] latex2exp_0.5.0     tools_4.1.0         backports_1.2.1
## [10] utf8_1.2.2          R6_2.5.0            DBI_1.1.1
## [13] colorspace_2.0-2    withr_2.4.2         tidyselect_1.1.1
## [16] gridExtra_2.3       prettyunits_1.1.1   processx_3.5.2
## [19] bit_4.0.4           curl_4.3.2          compiler_4.1.0
## [22] cli_3.0.1           rvest_1.0.1         xml2_1.3.2
## [25] labeling_0.4.2      scales_1.1.1        callr_3.7.0
## [28] digest_0.6.27       StanHeaders_2.21.0-7 rmarkdown_2.9
## [31] pkgconfig_2.0.3     htmltools_0.5.1.1   dbplyr_2.1.1
## [34] rlang_0.4.11        readxl_1.3.1        rstudioapi_0.13
## [37] generics_0.1.0      farver_2.1.0        jsonlite_1.7.2
## [40] vroom_1.5.3         distributional_0.2.2 inline_0.3.19
## [43] loo_2.4.1           Rcpp_1.0.7          munsell_0.5.0
## [46] fansi_0.5.0         viridis_0.6.1       lifecycle_1.0.0
## [49] stringi_1.7.3       yaml_2.2.1          ggraph_2.0.5
## [52] MASS_7.3-54         pkgbuild_1.2.0      parallel_4.1.0
## [55] ggrepel_0.9.1       crayon_1.4.1        lattice_0.20-44
## [58] graphlayouts_0.7.1  haven_2.4.3         hms_1.1.0
## [61] knitr_1.33          ps_1.6.0            pillar_1.6.2
## [64] igraph_1.2.6        codetools_0.2-18    stats4_4.1.0
## [67] reprex_2.0.1        glue_1.4.2          evaluate_0.14
## [70] V8_3.4.2            RcppParallel_5.1.4  modelr_0.1.8
## [73] vctrs_0.3.8         tzdb_0.1.2          tweenr_1.0.2
## [76] cellranger_1.1.0    gtable_0.3.0        polyclip_1.10-0
## [79] assertthat_0.2.1    xfun_0.24           ggforce_0.3.3
## [82] broom_0.7.9         tidygraph_1.2.0     coda_0.19-4
## [85] viridisLite_0.4.0   ellipsis_0.3.2
```
